# Harnessing the power of whole human liver ex situ normothermic perfusion for preclinical AAV vector evaluation

**DOI:** 10.1101/2023.07.10.548299

**Authors:** Marti Cabanes-Creus, Sophia H.Y. Liao, Renina Gale Navarro, Maddison Knight, Deborah Nazareth, Ngee-Soon Lau, Mark Ly, Erhua Zhu, Ramon Roca-Pinilla, Ricardo Bugallo Delgado, Grober Baltazar, Adrian Westhaus, Jessica Merjane, Michael Crawford, Geoffrey W. McCaughan, Carmen Unzu, Gloria González-Aseguinolaza, Ian E. Alexander, Carlo Pulitano, Leszek Lisowski

## Abstract

Developing clinically predictive model systems for evaluating gene transfer and gene editing technologies has become increasingly important in the era of personalized medicine. Liver-directed gene therapies present a unique challenge due to the complexity of the human liver. In this work, we describe the application of whole human liver explants in an *ex situ* normothermic perfusion system to evaluate a set of fourteen natural and bioengineered adeno-associated viral (AAV) vectors directly in human liver, in the presence and absence of neutralizing human sera. Under non-neutralizing conditions, the recently developed AAV variants, AAV-SYD12 and AAV-LK03, emerged as the most functional variants in terms of cellular uptake and transgene expression. However, when assessed in the presence of human plasma containing anti-AAV neutralizing antibodies (NAbs), vectors of human origin, specifically those derived from AAV2/AAV3b, were extensively neutralized, whereas AAV8-derived variants performed efficiently. This study establishes the use of normothermic liver perfusion as an invaluable preclinical model for evaluating liver-targeted gene therapies and providing guidance for making essential decisions that promote the most effective translational programs.

## Introduction

As we venture further into the era of personalized medicine, the challenge of developing clinically predictive model systems for evaluating gene transfer and gene editing technologies presents a critical hurdle. The ideal model system should not only accurately mimic human physiological conditions and intricate tissue organization, but also reliably predict clinical outcomes. Within the world of liver-directed gene therapies, 2D and 3D *in vitro* models, including induced pluripotent stem cell (iPSC) derived organoids, murine models including xenograft models, and non-human primates have traditionally been employed to develop and validate novel advanced therapeutics. However, given the fact that the liver is a complex organ composed of multiple cell types that contribute to unique structural and functional organization, the complexity of the human liver has proven difficult to recapitulate. In response to this issue, particularly concerning adeno-associated viral (AAV) vector-based liver-targeted therapies, we herein present the development and validation of a novel preclinical model based on whole human liver explants maintained in an *ex situ* normothermic (36°C) perfusion system. This model provides a platform that allowed us, for the first time, to assess the functionality of existing and novel recombinant adeno-associated viral (rAAV) vectors directly in the entirety of the human liver.

It is noteworthy that AAVs have recently attracted attention as effective and clinically proven gene therapy vectors. To date, five serotypes (AAV1, AAV2, AAV5, AAV9, and AAV-rh74) have earned regulatory approval for use in human patients^1^, including one marketed product for the treatment of haemophilia (Hemgenix™, based on AAV5) and most recently another for Duchenne Muscular Dystrophy (based on AAVrh74). However, the current generation of AAVs remain suboptimal for the majority of clinical applications, often requiring the use of high vector doses.^2^ Notably, prior to clinical evaluation, these AAV variants have been functionally tested in animal models, which while useful to evaluate safety and biodistribution, do not adequately recapitulate the intricate cellular dynamics and physiological responses of human livers.

To address this limitation, we turned our attention to normothermic human liver perfusion. The normothermic maintenance of the organ has been shown to reduce preservation-related graft injury compared to static cold storage in transplantable livers^3^, thus offering a potentially superior model for evaluating systemic delivery and the complex interactions between AAV, the intravascular compartment, egress to the liver and primary parenchymal and non-parenchymal human liver cells. Using the currently available protocols, livers are supplied with oxygen and nutrients at physiological temperature and pressures, maintaining conditions that support homeostasis, normal metabolic activity, and objective assessment of function in real-time.^3^ More recently, *ex situ* normothermic perfusion has also been evaluated as a method to enable liver splitting before transplantation, aiming to help to address donor shortages by facilitating the transplant of one pediatric and one adult recipient from a single donor.^4^ From the perspective of preclinical model for evaluation of advanced therapeutics, *ex situ* liver splitting also provides a promising model for evaluating gene therapy vectors and liver-directed biotherapeutics with a genetically matched control.^4^

In this work, we used two whole human liver explants perfused with human blood to perform a functional evaluation of natural and bioengineered AAV vectors in the presence or absence of neutralizing antibodies. Specifically, we performed a next-generation sequencing (NGS) based parallel comparison of the transduction profile of a set of fourteen AAV variants (**Supplementary Table 1**) in whole human livers that were deemed unsuitable for transplantation. To provide preclinical context, we compared, in parallel, the same vector mix in other commonly used liver preclinical models, namely in the murine model, in xenografted mice engrafted with primary human or non-human primate hepatocytes, and in non-human primates.

Together, our work broadens the repertoire of preclinical models available for conducting liver-directed vector studies. In the future, this model could also be evaluated for studies aimed at developing novel bioengineered AAV variants and for disease-specific phenotype correction using gene addition and editing approaches. Concurrently this would allow the study of critical translational parameters such as therapeutic vector doses and potential vector-induced toxicity in perhaps what is the closest preclinical model of the human liver to date - the human liver itself.

## Results

### Study design

To enable functional evaluation of AAV variants in the context of the whole human liver, we adapted the Liver Assist perfusion system to allow for prolonged perfusion times of up to a week.^5^ This system uses an open venous reservoir and incorporates two long-term oxygenators, a gas blender equipped with a pediatric flow regulator for ventilation control, and a flow-adjustable dialysis membrane for water-soluble toxin filtration and perfusate volume control (**Fig. 1a**).

**Figure 1.**
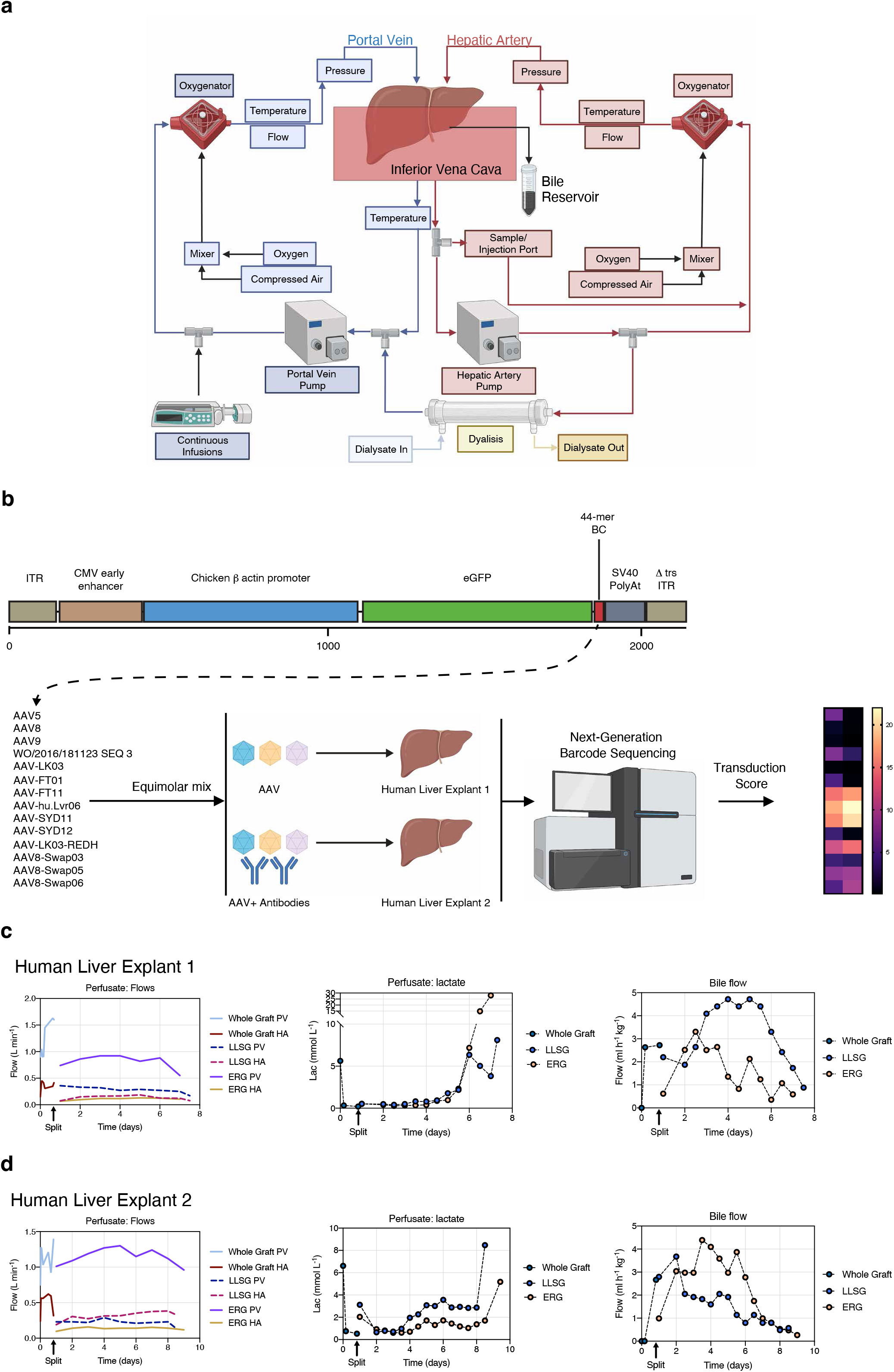
Study design and liver performance parameters during *ex situ* normothermic perfusion. **a,** Graphical flow chart of the liver perfusion machine. **b,** Transgene composition and study design. CMV - Cytomegalovirus; ITR – Inverted Terminal Repeat. **c,** Course of flows, lactate concentration and bile flow throughout the perfusion of Donor 1. **d,** Similar course analyses for Donor 2. PV – Portal Vein; HA – Hepatic Artery; LLSG - Left Lateral Sector Graft; ERG – Extended Right Graft.

To better recapitulate physiological conditions, we used a blood-based leukocyte-free perfusate, supplemented with essential nutritional supplements including amino acids, lipids, insulin, glucagon, taurocholic acid, and methylprednisolone (**Materials and Methods**).^4^ To minimize interference with AAVs that utilize heparan sulfate proteoglycan (HSPG) for cellular attachment, we supplemented the perfusate with enoxaparin (Clexane™), instead of standard heparin, as an anticoagulant. Before commencing the studies, we pre-screened numerous lots of fresh frozen plasma (FFP) for the presence of anti-AAV antibodies. This prescreening allowed us to perfuse two livers differently; one with a perfusate containing AAV Nabs-positive plasma (creating neutralizing conditions) and the other with a perfusate containing plasma with low levels of NAbs (creating non-neutralizing conditions) (**Fig. 1b**).

To facilitate multiplexed vector comparison, we validated a collection of self-complementary AAV (scAAV) transgenes encoding a CAG-eGFP-BC-pA expression cassette containing a unique 44-mer barcode (BC) between the eGFP reporter and the poly(A) (**Fig. 1b, Supplementary Fig. 1**). Each of the uniquely barcoded constructs was packaged into each of the fourteen selected AAV variants (**Fig. 1b** and **Supplementary Table 1)**. The selection included variants previously evaluated in clinical studies targeting the liver, such as AAV5, AAV8, and AAV-LK03, and also encompassed capsids used in clinical studies for organs other than the liver following systemic administration, such as AAV8 and AAV9. We also included several next-generation bioengineered capsids, such as AAV-SYD11 and AAV-SYD12^6^, which showed superior efficiency at transduction of primary human hepatocytes in other pre-clinical models of human liver. Each vector preparation was tittered individually, combined at a 1:1 molar ratio, and the composition of the transgene mix validated using NGS (**Supplementary Fig. 2**).

### Liver grafts, splitting and prolonged *ex situ* normothermic perfusion

Two whole human liver explants referred to as donor 1 (a 1.65 kg liver from a 78-year-old female) and donor 2 (a 2.19 kg liver from a 56-year-old male) were obtained from DonateLife, the centralized donation organization in Australia. These livers were unsuitable for transplantation and consented for research use. The first liver was excluded from transplantation due to biliary sepsis and cholecystostomy, while the second liver was donated after circulatory death (DCDD) and was deemed ineligible for transplantation due to the donor’s age (>50), though it was otherwise healthy. The livers were procured as detailed in the **Materials and Methods** section. Following procurement, the livers were transported in static cold storage solution under ischemic conditions and remained in these conditions for 187 and 305 minutes, respectively, from the time of retrieval until the start of normothermic perfusion.

Both liver explants met viability-based inclusion criteria (outlined at the **Materials and Methods section**) at the onset of the perfusion and again 24 hours later. Each liver was then split into two grafts: a left lateral sector graft (LLSG, consisting of segments 2 and 3) and an extended right graft (ERG, consisting of segments 1 and 4-8), following a recently described method.^4^ These partial grafts were then perfused concurrently, each with its own modified Liver Assist perfusion system.

For donor 1, we achieved a portal flow rate of 1.6 L min^−1^ and a pulsatile flow rate of 0.41 L min^−1^ for the hepatic artery. After splitting the organ, the flow rates were adjusted in accordance with the vessel split; the main portal vein flow for the extended right graft and the main hepatic artery flow for the left lateral sector graft. This liver exhibited immediate lactate clearance and progressively increasing bile production, which remained stable until day six before starting to decline (**Fig. 1c**). We terminated the experiment at one week from the initiation of the perfusion. Liver function, as gauged by Factor V synthesis, improved from an initial 13%, peaking at 29% for the LLSG 24 hours post-split and at 50% for the ERG 48 hours post-split (**Supplementary Fig. 3**). The liver transaminases release was high, as we recorded Alanine aminotransferase (ALT) at 522 U L^−1^ at 24 hours post-split for the LLSG and 2,105 U L^−1^ for the ERG at the same time point. The ALT levels continued to rise throughout the perfusion, indicating likely reperfusion injury (**Supplementary Fig. 4**). We noted a similar trend with the acute cytokine levels in the perfusate, which began at a relatively low level (IL-6 = 78.8 ng L^−1^, 4 hours post-perfusion), and spiked to over 10,000 ng L^−1^ for both grafts by day five post-splitting (**Supplementary Fig. 5**).

In the case of donor 2, we observed similar trends in perfusate flows, lactate clearance, and bile production, but this liver remained stable for a slightly longer duration (**Fig. 1d**). The functionality of this liver was reflected by the rise in Factor V synthesis from 14% at the beginning of perfusion, reaching a peak of 102% for the LLSG and 105% for the ERG five days post-split (**Supplementary Fig. 6**). The ALT levels peaked at 9,762 U L^−1^ at 4 hours post-split for the LLSG, and gradually decreased to 1,663 U L^−1^ on day 7.5 after splitting (**Supplementary Fig. 7**). For the ERG, ALT levels rapidly increased, peaking at 6,516 U L^−1^, 4 days post-split, and then declining to 2,273 U L^−1^ on day 8 post-split (**Supplementary Fig. 7**). The acute cytokine IL-6 levels in the perfusate stayed relatively stable until day 7, at which point they rapidly increased to 1,622 ng L^−^ for the left graft and at 15,744 ng L^−^ for the right graft (**Supplementary Fig. 8**). Overall, these results suggest that both livers were stable for a period of up to six days.

### Pre-screening of fresh frozen plasma for the presence of anti-AAV antibodies

In order to compare the efficiency of the chosen AAV variants (**Supplementary Table 1**) both with and without the interference of anti-AAV neutralizing antibodies (Nabs) (**Fig. 1b**), we first conducted a pre-screening of eighteen lots of human plasma. Relying on the established correlation between ELISA-based and neutralization-based assays^7^, we followed a recently described ELISA procedure^8^ to examine each plasma against the equimolar mixture of the fourteen AAV vectors. We classified the plasma samples based on their total reactivity (**Supplementary Fig. 9**).

Following preparation of the final plasma-containing perfusate, we examined the reactivity of each perfusate against each of the individual capsids present in the AAV mix. To do so, as outlined in the **Materials and Methods** section, we evaluated the reactivity of each of the 14 capsid variants with the perfusate at dilutions of 1:25, 1:50, and 1:100. With the non-reactive perfusate used for donor 1 (**Supplementary Fig. 10**), we found all capsids to have an end titer of <1:25, excluding AAV9, which showed minor reactivity at this dilution, and is thus reported as presenting an end titer of 1:25. For the reactive perfusate used with donor 2, we found end titers to be 1:50 for the AAV3b-based variants (AAV-LK03, AAV-LK03-REDH, and AAV-SEQ-3) and for AAV-FT11, 1:25 for AAV-FT01, AAV-hu.Lvr06, AAV-SYD11, and AAV-SYD12, and non-reactive or <1:25 for AAV5, AAV8, AAV9, AAV8-Swap03, AAV8-Swap05, and AAV8-Swap06 (**Supplementary Fig. 11**).

### Functional evaluation of AAV vectors in the *ex situ* human liver perfused with non-neutralizing human plasma

With the appropriate human plasma samples identified, we proceeded to perfuse the first liver under non-neutralizing conditions. As schematically depicted in **Fig. 2a**, seven hours after starting the perfusion, and once the lactate levels had dropped from 5.62 mmol L^−1^ to 0.33 mmol L^−1^ (**Fig. 1c**), we injected the vector mix to the whole graft prior to liver splitting. We administered a total of 3.10×10^12^ vector genomes (vg) directly into the portal vein. This dosage corresponds to 1.92×10^12^ vg per kg of liver, which would equate to an estimated dose of 4.5×10^10^ vg per kg of body weight. Considering the mix contained fourteen capsids, the estimated dose per capsid was 1.37×10^11^ vg per kg of liver. Approximately fourteen hours post-AAV injection, we split the liver into the left lateral sector graft (LLSG) and the extended right graft (ERG), while maintaining uninterrupted arterial and portovenous perfusion (**Materials and Methods).** Subsequently, we independently perfused and consistently monitored both grafts using separate Liver Assist machines. Notably, the LLSG continued to be perfused with the original perfusate containing AAVs, while the ERG received new AAV-free perfusate. This setup allowed us to investigate the kinetics of transduction at the fourteen-hour mark by comparing the AAV transduction profiles in both liver grafts (**Fig. 2a**).

**Figure 2.**
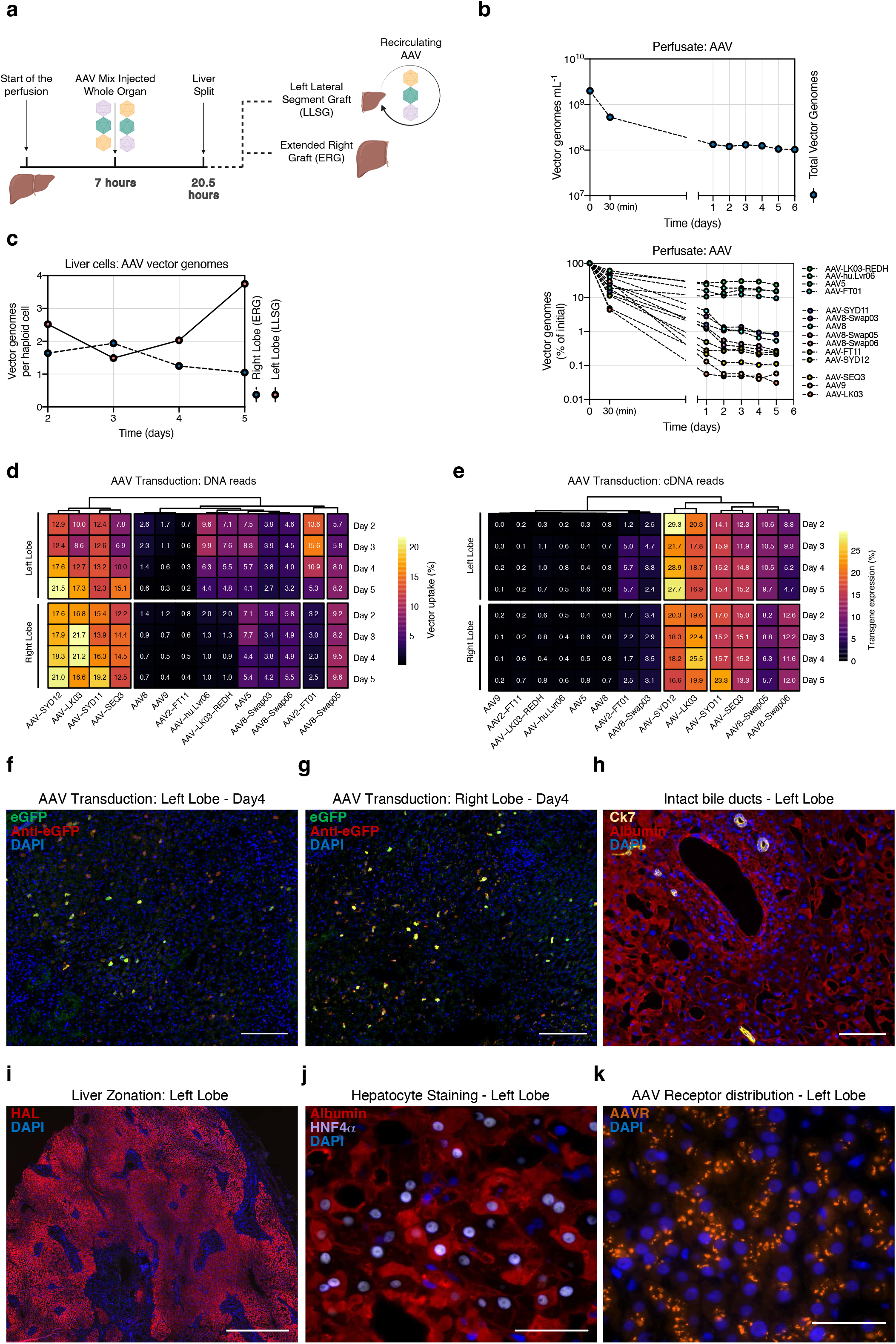
Functional evaluation of AAV vectors in the *ex situ* human liver perfused with non-neutralizing human plasma. **a,** Graphical representation of the study. **b,** Concentration of total (top panel) and de-multiplexed AAV vector genomes in the perfusate (bottom panel). Concentration of individual vectors is expressed as the estimated percentage of the initial concentration as studied with NGS. Vectors are ranked from most to least abundant (top to bottom). **c**, Total vector genomes per haploid cell found in biopsies from both liver grafts extracted at the indicated perfusion times. **d,** Percentage of NGS reads mapped to each barcoded AAV capsid variant. The transgene DNA, indicating vector uptake, was extracted from biopsies taken from both grafts at the indicated time points. **e,** Similar analysis performed on transgenes recovered from RNA, which indicate functional transduction. Percentages are normalized to the pre-injection mix. **f,** Representative immunofluorescence analysis of the net eGFP signal from collective AAV transduction in the left graft. **g,** Similar representative immunofluorescence image for the right graft. The vector-encoded eGFP was also counterstained with an anti-eGFP antibody (red). Blue: DAPI (nuclei). Scale = 100 μm. **h,** Representative immunofluorescence analysis of the bile ducts and hepatocytes of the left graft. Yellow: Cytokeratin 7 (bile ducts); red: albumin; blue: DAPI (nuclei). Scale: 50 μm. **i,** Representative immunofluorescence analysis of liver zonation. Red: Histidine ammonia-lyase (HAL); blue: DAPI (nuclei). Scale: 500 μm. **j,** Representative immunofluorescence analysis of liver integrity. Red: albumin; Purple: Hepatocyte nuclear factor 4 alpha (HNF4α); blue: DAPI (non-hepatocyte nuclei). Scale: 20 μm. **h,** Representative immunofluorescence analysis of AAV Receptor (AAVR) in the human liver. Orange: AAVR; Blue: DAPI (nuclei). Scale: 100 μm.

We monitored vector clearance from the perfusate by analyzing plasma samples for the presence of AAVs at various times after vector infusion (**Fig. 2b**). Quantification of total AAV load indicated that starting from the initial concentration of 1.55×10^9^ vg mL^−1^, the concentration of AAVs declined sharply to approximately 9% of these initial levels 24 hours post-injection (**Fig. 2b**). Subsequently, the AAV levels in the perfusate stabilized, suggesting that after the target organ absorbed the initial vector dosage, a noticeable fraction of around 10% of the vector mix persisted in circulation for the rest of the study.

To identify which of the AAV variants from the infused mix exhibited the lowest rates of clearance in the perfusate we performed NGS quantification of the unique barcoded region of the transgenes recovered from the perfusate, thereby demultiplexing the fraction of each AAV. In evaluating the percentage drop of each individual AAV over time in the perfusate, we distinguished three vector groups (**Fig. 2b**). A group of four variants - the HSPG de-targeted AAV-LKO3-REDH, AAV-hu.Lvr06, AAV5, and AAV-FT01, which were the primary contributors of the vectors present in the perfusate after the initial vector clearance (**Fig. 2b**). Following the initial drop, these variants persisted in the circulation at levels above 10% of the initial concentration. Another group, namely AAV9 and the HSPG-binders (AAV-LK03 and AAV-SEQ3), demonstrated the fastest clearance kinetics from the perfusate (**Fig. 2b**), with vector levels dropping to approximately 0.1% or less of the initial vector concentration within two days. All the remaining AAV variants exhibited an intermediate phenotype (**Fig. 2b**), with final concentrations varying between 0.1% and 1% of the individual initial levels.

We then turned our focus to the overall transduction profile of the vector mix in both liver grafts. To do this, we evaluated the total vector copy number per haploid cell in the DNA extracted from liver biopsies on days two to five post-injection. We found an increase in the average vector copy number in the left graft (LLSG) that continued to be perfused with the AAV-containing perfusate following the organ split (**Fig. 2c**). In contrast, the vector copy number in the right graft, which received AAV-free perfusate after the organ split, remained relatively stable at the tested timepoints, ranging from 1.5 to 1 vg per haploid genome (**Fig. 2c**).

To further analyze the transduction profile, we used NGS to examine the barcode composition in the DNA and RNA samples (which are indicative of cell entry performance and transgene expression, respectively) extracted from liver biopsy samples. We first calculated the percentage contribution of each vector to the total vector genomes in each graft, normalizing the data to the input pre-injection mix (**Supplementary Fig. 2**). We then used unsupervised hierarchical clustering to group the AAV vectors with similar transduction profiles in the human liver, both at the DNA (vector uptake, **Fig. 2d**) and the RNA (functional transduction, **Fig. 2e**) levels.

As previously discussed, the right extended graft (ERG) provided a unique opportunity to study vector uptake during the first 14 hours post-injection. The AAV-SYDs and AAV-LK03 displayed rapid transduction kinetics, closely followed by the AAV3b-based AAV-SEQ3. In contrast, we found that AAV8, AAV9, AAV-FT1 and AAV-F11, AAV-hu.Lvr06, and AAV-LK03-REDH sat at the other end of the spectrum. These variants either did not efficiently transduce this liver or demonstrated slower kinetics of vector uptake (**Fig. 2d**, bottom panel). Looking at transduction kinetics throughout the experiment, we found that the overall vector transduction performance remained stable throughout the experiment (**Fig. 2d**, bottom panel). The transduction data we obtained from the left graft (LLSG) appeared more variable than the data for the right graft (refer to top panel of **Fig. 2d**). We believe this was likely a consequence of vector recirculation in the perfusate following organ split, which facilitated ongoing transduction during the course of the study. Specifically, in the LLSG, we detected increased relative transduction of some variants over time (AAV-SYD12, AAV-LK03, and AAV-SEQ3), while others exhibited decreased relative transduction over time (AAV-FT01, AAV-hu.Lvr06, AAV-LK03-REDH). Transduction for the remaining vectors remained stable.

Since these data present relative transduction, a decreasing contribution in the liver biopsies likely indicates either slower transduction kinetics or a decreasing amount of artificial signal stemming from the interstitial perfusate, rather than from vector uptake in hepatocytes. Indeed, the vectors showing a decreased relative transduction over time are the same one that stayed longer in the perfusate (**Fig. 2c**).

One interesting cluster of vectors comprised AAV-FT01, AAV-hu.Lvr06 and AAV-LK03-REDH. These vectors seemed to transduce the left graft but failed to transduce the right graft, which was exposed to the vectors for only the first fourteen hours. This likely indicates a slower transduction kinetics of these vectors (**Fig. 2d**).

Subsequently, we evaluated the barcode composition at the cDNA level in both grafts, serving as an indicator of each variant’s efficiency at functionally transducing cells in this preclinical model of the human liver (**Fig. 2e**). Again, we employed unsupervised hierarchical clustering of the data to identify variants presenting similar functional transduction. Variants like AAV5, AAV-hu.Lvr06 and AAV-LK03-REDH did not functionally transduce the liver, despite the fact that these vectors were detected at the DNA level (**Fig. 2e**). Conversely, other variants like the AAV-SYDs, AAV-LK03, AAV-SEQ3, and AAV8-Swap05 (a variant harboring variable region (VR-I) from AAV2 and VRs VI to VIII from AAV7) exhibited similar functional transduction kinetic, with levels of reads recovered from cDNA closely matching the levels recovered from DNA. The AAV8-Swap06 variant displayed faster functional transduction kinetics, evident by a higher relative percentage of cDNA reads (~2×) compared to DNA reads at all the studied time points, a phenomenon we had previously observed in mice with humanized livers.^6^

Lastly, we conducted microscopic analysis of the transduced liver using immunofluorescence (**Fig. 2f-j**) as well as hematoxylin and eosin staining (H&E, **Supplementary Fig. 12**). We could readily detect eGFP signal arising from collective vector transduction at day 4 post-injection in both grafts (**Fig. 2f-g, Supplementary Figs 13-14**). Notably, intrahepatic bile ducts appeared well-preserved (**Fig. 2h**), as was liver zonation, demonstrated by staining of the portal marker histidine ammonia-lyase (HAL, **Fig. 2i**). We confirmed liver parenchymal and non-parenchymal integrity through staining for albumin and the hepatocyte nuclear factor 4 (HNF4A), which stains for human hepatocytes nuclei (**Fig. 2j**). Finally, to gain insights into the mechanisms of cell-AAV interaction, we analyzed the distribution of the AAV-Receptor (AAVR) in the human liver explant (**Fig. 2k**). AAVR could be detected in human hepatocytes, although the localization of appeared to be primarily intracellular (**Fig. 2k**).

### Functional evaluation of AAV vectors in the *ex situ* human liver perfused with neutralizing human plasma

In the second experiment, we perfused the donor 2 liver using plasma containing antibodies reactive against a subset of capsids present in the vector mix (**Supplementary Fig. 11**). Unlike the first study detailed earlier, we only injected the vector mix into the left graft (**Fig. 3a**), post-liver split, once the lactate level had reduced from 3.12 mmol L^−1^ to 0.65 mmol L^−1^ (**Fig. 1d**). The right graft did not receive any AAV and acted as a donor-matched, untreated control. In addition, to counter the presence of neutralizing antibodies for some of the capsids, we doubled the dose of AAV per kg of liver relative to the first liver. Specifically, we injected a total of 2.57×10^12^ vector genomes into the left portal vein. Given that the LLSG weighted 0.67 kg, this corresponded to a total dose of 3.83×10^12^ vg per kg of liver, or 2.74×10^11^ vg per kg of liver for individual variants present in the mix.

**Figure 3.**
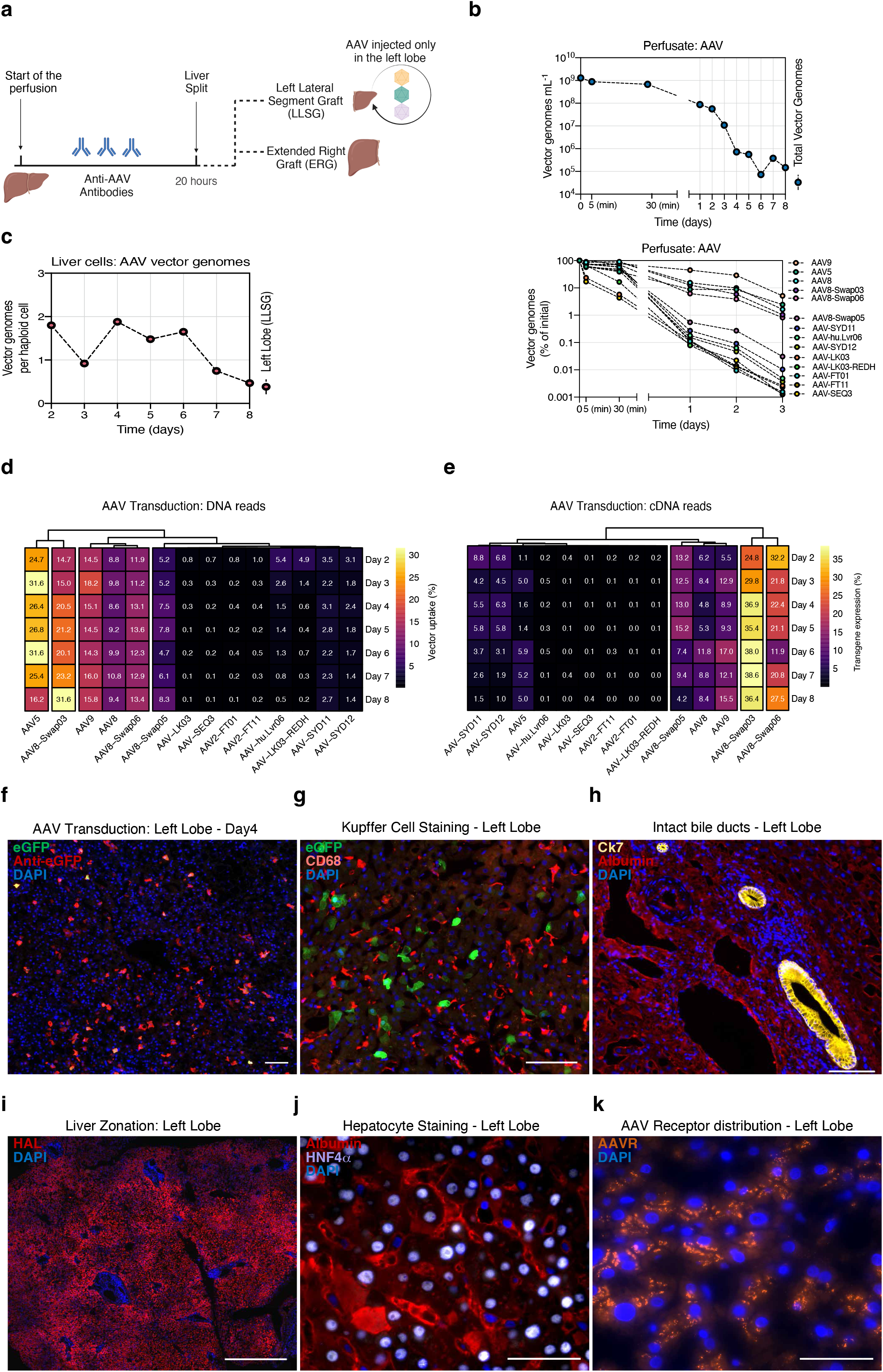
Functional evaluation of AAV vectors in the *ex situ* human liver perfused with neutralizing human plasma. **a,** Graphical representation of the study. **b,** Concentration of total (top panel) and de-multiplexed AAV vector genomes in the perfusate (bottom panel). Concentration of individual vectors is expressed as the estimated percentage of the initial concentration as studied with NGS. Vectors are ranked from most to least abundant (top to bottom). **c**, Total vector genomes per haploid cell found in biopsies the left graft. **d,** Percentage of NGS reads mapped to each barcoded AAV capsid variant. The transgene DNA, indicating vector uptake, was extracted from biopsies taken from both grafts at the indicated time points. **e,** Similar analysis performed on transgenes recovered from RNA, which indicate functional transduction. Percentages are normalized to the pre-injection mix. **f,** Representative immunofluorescence analysis of the net eGFP signal from collective AAV transduction in the left graft. The vector-encoded eGFP was also counterstained with an anti-eGFP antibody (red). Blue: DAPI (nuclei). Scale = 100 μm. **g,** Representative immunofluorescence analysis of the presence of Kupffer cells in the left graft. Green: vector-encoded eGFP; Red: CD68 (Human macrophage marker); blue: DAPI (nuclei). Scale 100 μm. **h,** Representative immunofluorescence analysis of the bile ducts and hepatocytes of the left graft. Yellow: Cytokeratin 7 (bile ducts); red: albumin; blue: DAPI (nuclei). Scale: 50 μm. **i,** Representative immunofluorescence analysis of liver zonation. Red: Histidine ammonia-lyase (HAL); blue: DAPI (nuclei). Scale: 500 μm. **j,** Representative immunofluorescence analysis of liver integrity. Red: albumin; Purple: Hepatocyte nuclear factor 4 alpha (HNF4α); blue: DAPI (non-hepatocyte nuclei). Scale: 20 μm. **h,** Representative immunofluorescence analysis of AAV Receptor (AAVR) in the human liver. Orange: AAVR; Blue: DAPI (nuclei). Scale: 100 μm.

As in the case of donor 1, we initially measured vector clearance from the AAV-containing perfusate. In stark contrast to our observations under non-neutralizing conditions, where vector genomes stabilized at around 1×10^8^ vg mL^−1^ (**Fig. 2b**), we observed a rapid disappearance of vector genomes from the perfusate to below 1×10^6^ vg mL^−1^ by day 4 (**Fig. 3b**), equivalent to ~0.06% of the estimated initial concentration of 1.29×10^9^ vg mL^−1^. Next, we employed NGS to investigate the clearance kinetics of individual variants from the perfusate. Given the time-course of vector clearance (**Fig. 3b**), we focused on the changes in the composition of the AAV mix present in the perfusate during the first three days of the study. We identified two distinct vector populations; i) a group of five variants (AAV9, AAV5, AAV8, AAV8-Swap03 and AAV8-Swap06) that exhibited slow clearance kinetics, with each variant present at levels above 1% of the initial vector composition on day 3 post-injection (**Fig. 3b**, lower panel) and ii) a group consisting all remaining variants that were rapidly cleared from the system, with levels ranging from 0.001% to 0.01% of their corresponding initial levels, one-log lower than the lowest levels previously observed under non-neutralizing conditions (**Fig. 2b**). Notably, there appeared to be a correlation between high plasma reactivity and faster vector clearance kinetics from the perfusate, as all vectors present on day 3 at <1% of the initial concentration displayed an end Nab titer of 1:25 or >1:50 (**Supplementary Fig. 11**). This could suggest that the anti-AAV neutralizing antibodies present in the perfusate were directly responsible for vector removal from the circulation, most likely through opsonization followed by residential macrophage clearance. In fact, we found a statistically significant Spearman negative correlation between plasma reactivity and perfusate clearance (Day 1: r = −0.9165, p<2.2×10^−16^; Day 2: r = −0.8725, p<2.2×10^−16^; Day 3: r = −0.8901, p<2.2×10^−16^), in support of this hypothesis.

Analysis of the vector copy number in the biopsy samples taken between day 2 and day 6 post vector infusion indicated an average of ~1.5 vector genomes per haploid cell up to day six of the study, followed by a small yet noticeable decrease to around 0.5 vector genomes per haploid cell at the conclusion of the study on day 7 and 8 (**Fig. 3c**). Next, we used NGS to examine the transduction profile of individual AAVs in the left graft. **Fig. 3d** shows the hierarchical clustering of vector uptake (at the DNA level) contribution after normalization to the pre-injection input mix (**Supplementary Fig. 2**). In marked contrast with the non-neutralizing conditions for donor 1 (**Fig. 2d**), we did not detect AAV-LK03 uptake for this donor, and only a relatively low number of reads mapped to AAV-SYD12 (**Fig. 3d**). Generally, the presence of neutralizing antibodies for specific capsids effectively obstructed liver uptake of those variants (**Fig. 3d**). The only capsids we could readily detect in this liver were those that did not react with this batch of human plasma, namely AAV5, AAV8, AAV9, AAV8-Swap03 and AAV8-Swap06 (**Supplementary Fig. 11**). AAV8-Swap05, which displayed the slowest kinetics of clearance from perfusate among the group of reactive variants (**Fig. 3b**, lower panel), also exhibited a slightly higher antibody reactivity at 1:25 (**Supplementary Fig. 11**). Interestingly, AAV8-Swap03, which contains variable regions VI to VIII from AAV7,^6^ functioned particularly well under these neutralizing conditions. In terms of functional transduction, or cDNA reads, we noticed a trend similar to that observed for vector uptake. The sole exception was AAV5, which functioned relatively less efficiently than the other variants, especially considering its relatively higher vector uptake (**Fig. 3e**).

In the subsequent analysis of this liver using immunofluorescence (**Fig. 3f-j**) and H&E staining (**Supplementary Fig. 15**), we observed a similar pattern to that of donor 1. On day 4 post-injection, we could readily detect the net eGFP signal from collective transduction by the vectors present in the mix (**Fig. 3f**). Considering the fact that the vectors reactive with the anti-AAV Nabs present in the plasma were cleared from this closed system (we could not detect them either in the DNA extracted from the biopsy samples or within the perfusate), we investigated whether Kupffer cells (resident liver macrophages) were responsible for this active clearance. As shown in **Fig. 3g** consistent with the fact that this was an entire human liver organ, we could detect a large number of Kupffer cells (CD68+) in the liver. To investigate whether AAV particles were actively being cleared by Kupffer cells after opsonization with reactive antibodies, we co-stained liver sections with the anti-AAV B1 antibody, which stains for linear capsid epitopes, but failed to observe any detectable co-localizing signal (data not shown). Notably, just as described for the liver from donor 1, histological analyses confirmed that the intrahepatic bile ducts were well-preserved (**Fig. 3h**), as was metabolic zonation (**Fig. 3i**), parenchymal integrity (**Fig. 3j**) and wide lobular expression of the AAV Receptor (**Fig. 3k**).

### Functional evaluation of AAV vectors in murine and non-human primate liver models

Finally, to place the results observed in the human liver explant in the context of other established preclinical models of human liver, we studied the relative performance of the same set of fourteen vectors (**Supplementary Table 1**) in the murine liver, the xenograft mouse models engrafted with human and non-human primate hepatocytes, as well as *in vivo* in a non-human primate (**Fig. 4a**).

**Figure 4.**
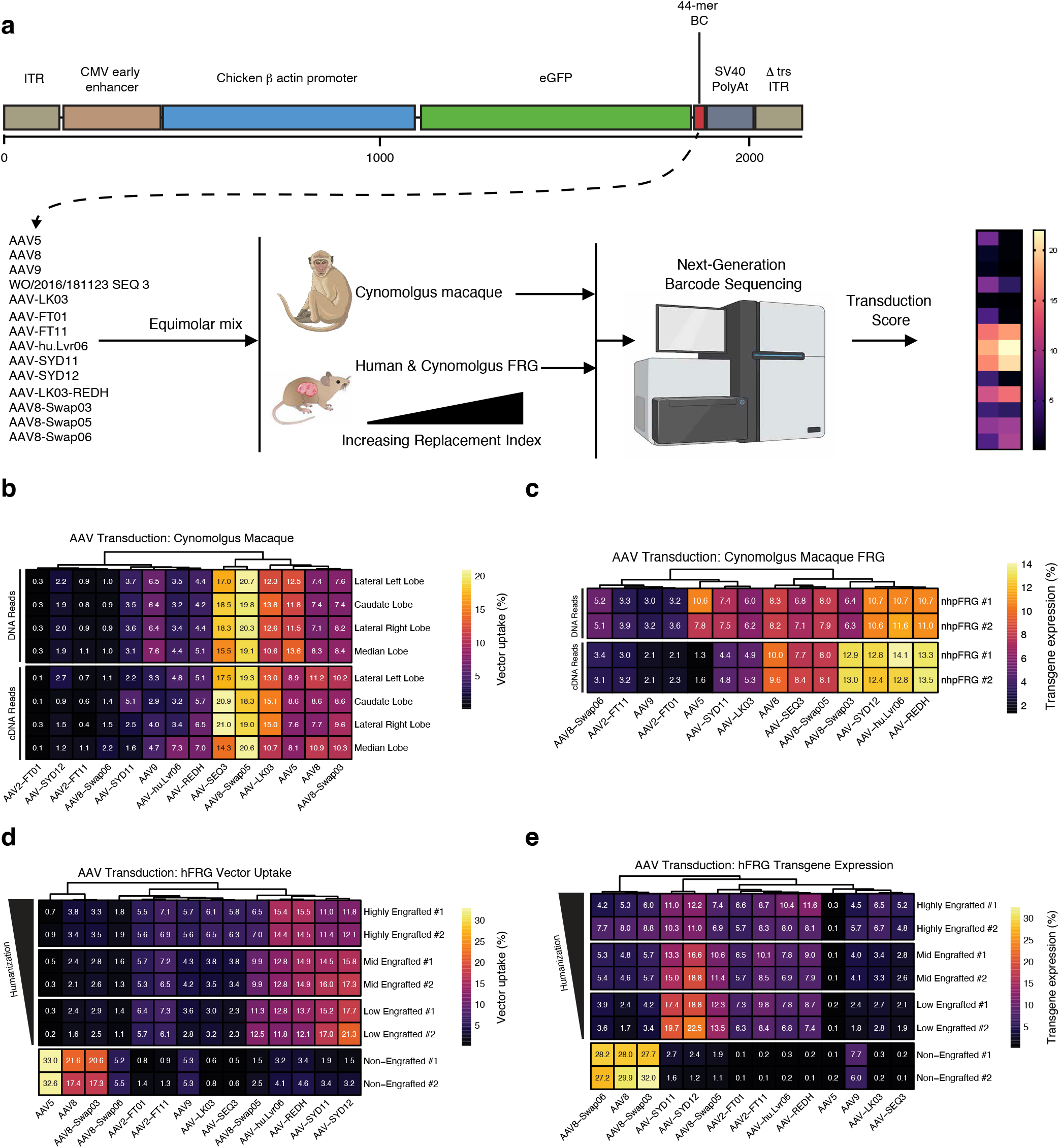
Functional evaluation of AAV vectors in murine and non-human primate liver models. **a,** Transgene composition and study design. CMV - Cytomegalovirus; ITR – Inverted Terminal Repeat. **b,** Percentage of NGS reads mapped to each barcoded AAV capsid variant. On the top panel, the transgene DNA, indicating vector uptake, was extracted from tissue taken from the four indicated lobes one-month post-injection. The bottom panel shows a similar analysis performed on transgenes recovered from RNA, which indicate functional transduction. Percentages are normalized to the pre-injection mix. **c,** Similar analyses performed on FRG mice engrafted with cynomolgus hepatocytes. The hepatocytes were sorted prior to analyses one-week post-injection. **d**, Similar analyses at the DNA level (vector uptake) performed in naïve FRG and FRG presenting different levels of humanization as indicated by the black scale. **e,** Similar analyses at the RNA/cDNA level.

First, we performed an *in vivo* evaluation of the same vector mix using the cynomolgus monkey NHP model (**Fig. 4b**). In order to minimize the influence of neutralizing antibodies, we removed the majority of antibodies using a previously validated immunoadsorption method.^9^ After antibody depletion, the end titer, calculated for AAV9, was 1:30 (**Materials and Methods**). We then intravenously infused a total dose of 3.5×10^12^ vector genomes (1.4×10^12^ vg kg^−1^) of a new barcoded AAV mix (**Supplementary Fig. 16**). One-month post injection, we analyzed the relative vector performance in the four liver lobes, both at the vector uptake (DNA) and transgene expression (mRNA/cDNA) levels using NGS of the unique barcoded region of the expression cassette. At this time point, the relative vector uptake was highly correlated with transgene expression (r = 0.96, p=4.98×10^−8^, Spearman’s correlation coefficient) (**Fig. 4b**). As seen previously in the human liver explant, the variants that did not follow this pattern were AAV5, which showed efficient uptake but low transgene expression (12.37% vs 8.31% average read share, respectively, for AAV5 **Fig. 4b**), and to a lower extent AAV9. The AAV8 variant AAV8-Swap05 (containing AAV2’s VR-I and AAV7’s VRs VI-VIII) performed more efficiently among all vectors tested, both at cell entry and transgene expression, followed closely by the AAV3b related AAV-SEQ3 and AAV-LK03. In marked contrast to the data obtained in human liver explants (**Fig. 2**), the AAV-SYDs failed to efficiently transduce liver of this NHP (**Fig. 4b**).

We next evaluated the vector mix in a recently developed FRG xenograft murine model engrafted with primary hepatocytes from cynomolgus monkeys.^9^ To do so, we injected highly engrafted animals (n=2) with albumin levels 10 mg mL^−1^ blood (equivalent to estimated >80% repopulation level) with 1×10^11^ total vg of the AAV mix (approximately 7.5×10^9^ vg per capsid) and harvested the chimeric livers two-weeks post-infusion.

The results from this model resembled those from the NHP only for AAV5 and AAV9, where we observed the same post-entry defect for both serotypes. However, the top-performing variants in the cynomolgus model (AAV8-Swap05, AAV-SEQ3, AAV-LK03) ranked as medium performers in the xenograft mice engrafted with cynomolgus hepatocytes. Meanwhile, the top-ranking variants in the FRG xenograft model (AAV-LK03-REDH, AAV-hu.Lvr06 and AAV-SYD12) performed relatively poorly in the NHP model *in vivo* (compare data in **Fig. 4b** and **4c**). These findings highlight important differences between the cynomolgus monkey NHP model and the xenograft murine model engrafted with primary hepatocytes from cynomolgus monkeys.

Next, we investigated the same set of variants in humanized FRG models with various replacement indices (low ~10%, medium ~40% and high >80%, n=2 per repopulation level), as well as naïve (non-engrafted) FRGs. To facilitate direct comparison, we maintained the vector dose at a constant 7.5×10^9^ vg per capsid (~1×10^11^ total vg) and harvested the mice one-week post vector injection. As depicted in relative AAV transduction in human hepatocytes displayed marked differences in comparison to the murine liver (non-engrafted animals) (**Fig. 4d** and **4e**). We found that AAV5, AAV8, and AAV-Swap03 efficiently transduced the murine liver (**Fig. 4d**), however, these same variants ranked poorly in human hepatocyte entry, irrespective of the level of engraftment (**Fig. 4d**).

Our detailed analysis further revealed intriguing trends regarding vector performance in this preclinical human liver model. As the repopulation level increased (human-to-murine hepatocytes ratio), the relative function of AAV-SYDs and AAV8-Swap05 decreased (**Fig. 4d**). Conversely, the relative function of AAV-hu.Lvr06, AAV-LK03-REDH and the HSPG-binders (AAV-LK03 and AAV-SEQ3) improved with an increase in the repopulation level (**Fig. 4d**). Overall, the AAV-SYDs, AAV-hu.Lvr06 and AAV-LK03-REDH performed the best in the humanized animals at vector uptake (**Fig. 4d**).

Regarding the transgene expression / functional transduction (RNA/cDNA), the results were even more pronounced in the murine liver, where only AAV8, AAV8-Swap03, and AAV8-Swap06 were able to achieve significant transgene expression (**Fig. 4e**). Consistent with the data obtained from other models, AAV5 showed poor transgene expression, even though it was highly efficient at entering murine hepatocytes (**Fig. 4d-e**).

Of note, in human hepatocytes, we also noted a relatively lower cDNA read share for AAV-LK03-REDH and AAV-hu.Lvr06 when contrasted to their performance at vector uptake, setting them apart, at least functionally, from the AAV-SYDs (**Fig. 4d vs 4e**). To further investigate this phenomenon, we conducted two additional studies in the humanised FRG model. In the first, we infused two highly engrafted animals with an identical total vector dose, but extended the experiment for two months before harvesting the hepatocytes for subsequent vector NGS analyses. In the second study we injected two highly engrafted animals, but in this case, we doubled the total vector dose, resulting in each animal receiving 1.5×10^10^ vg per capsid (~2.1×10^11^ total vg). We then perfused the livers and isolated the hepatocytes one-week post-injection. The results demonstrated that increasing the vector dose had a minimal effect on relative performance (**Supplementary Fig. 17**). Conversely, the correlation between cDNA and DNA was considerably higher at the two months’ time point (r = 0.94, p=1.8×10^−8^) than at one-week post injection (r = 0.79, p=8.0×10^−4^) (**Supplementary Fig. 18**). This suggests that the expression kinetics, the aggregate result of the vector’s entire intracellular journey, is an important and differing factor when evaluating relative vector function. Importantly, the correlation of DNA read share between both time points was also high (r = 0.92, p=2.7×10^−8^), further underlining the significance of considering the influence of vector trafficking kinetics on overall vector function.

## Discussion

In 1976, a British statistician George Box^10^ made a famous statement: “Remember that all models are wrong; the practical question is how wrong do they have to be to not be useful”. This has become an enduring reminder that we must constantly assess the efficacy of the models we employ in research. In this study, we pioneered the use of an *ex situ* whole human liver model that, while not free of limitations, provides an unprecedented platform to explore the intricate workings of AAV vector performance. The model, involving a human organ perfused with human blood and maintained at body temperature, offers a strikingly accurate approximation of the human physiological setting. One could argue that there are few alternatives that bring us closer to the human setting, short of an actual clinical trial. This signifies the extraordinary potential of this novel model in accelerating progress in the field of liver mediated gene therapies – both viral and potentially non-viral - by enabling investigations that closely mimic the clinical scenarios where gene therapies will ultimately be deployed.

As evidenced in our study, *ex situ* human liver explants can be maintained in a viable state over an extended period, thereby providing a sufficiently long experimental window to facilitate a broad range of studies, including the development and evaluation of viral vectors. In the perfusions detailed here, liver function remained stable up to the sixth day, as demonstrated by the steady perfusate flow, consistently low lactate concentration in the perfusate, and sustained bile production (**Fig. 1c**). Nonetheless, *a posteriori* analysis of the samples revealed a substantial release of alanine aminotransferase (ALT) (**Supplementary Figs 4 and 7**). Considering the fact that these increases were already evident prior to vector injection and in samples collected from the control untreated right graft of donor 2, which was not exposed to the AAV vector (**Fig. 3a**), it is reasonable to attribute the elevated ALT to reperfusion injury. Reperfusion injury is a significant challenge in liver transplantation^11^ and, in this instance, was likely unrelated to the vector transduction process. In our study, we incorporated the use of methylprednisolone to limit reperfusion injury and associated adverse effects. Nonetheless, future research is necessary to pinpoint the most effective treatment regimen.

One significant advantage of the liver explant system is the opportunity to investigate antibody neutralization in an environment containing liver-resident immune cells. Intriguingly, we noted a rapid and almost complete neutralization of AAV variants that displayed reactivity to neutralizing antibodies (NAbs) on end-point titers of 1:50 and above. This led to rapid clearance from the perfusate (**Fig. 3b**), an effect we hypothesize to be mediated by liver-resident Kupffer cells. It is noteworthy that this phenomenon would be challenging to replicate in any existing preclinical liver model. We observed minor reactivity of the overall non-reactive perfusate (from donor 1) to AAV9 (**Supplementary Fig. 10**), which exhibited rapid clearance from the perfusate under these conditions (**Fig. 2b**, lower panel). Consequently, it is reasonable to hypothesize that AAV9’s modest performance in donor 1 (**Fig. 2d-e**) could be due to antibody neutralization. Considering the fast clearance kinetics observed when the vector mix was directly injected into the portal vein, systemic administration of vectors may prove to be even more challenging. Interestingly, and perhaps predictably, given our use of plasma from human donors, who are more likely to possess antibodies against viruses infecting humans, the vectors that performed best under non-neutralizing conditions were also those most heavily neutralized. Conversely, the AAV vectors that managed to evade this specific human plasma performed relatively poorly under non-neutralizing conditions. This observation hints at a potential trade-off between function and antibody neutralization. These findings also reinforce the hypothesis that under neutralizing conditions, strategies aimed at circumventing anti-AAV antibodies, such as the employment of endopeptidases^12^, could prove critically important in enhancing the potential for patients with detectable anti-AAV NAbs to benefit from innovative gene therapies.

A second powerful feature of the human liver explant model is the potential it offers for aligning vector functionality with historical clinical data. In our study, we administered a relatively low vector dose to both livers, a quantity significantly smaller than those utilized in some clinical trials. For donor 1, the injection of the vector mix into the entire organ allowed for an estimation of the equivalent vector dose per kg of body weight, which amounted to approximately 4.5×10^10^ vg per kg of body mass, based on the known body mass of the donor of liver 1. As the mix contained 14 AAV variants, the calculated dose per capsid was only 3.2×10^9^ vg per kg of body weight. This represents a figure nearly 20,000 times lower than the dose administered in Roctavian, the recently AAV5-based product for Hemophilia A.^13^ Our inability to detect transgene expression at the studied time-points for AAV5 implies that a higher dose might be necessary for this specific serotype (**Fig. 2e**). However, the detection of eGFP expression under both neutralizing and non-neutralizing conditions, attributable to other liver-tropic capsids present in the mix, suggests that lower clinical vector doses may be sufficient to achieve clinical benefits if AAV variants that transduce human primary hepatocytes with high efficiency are used.

This study also allowed us to compare data obtained from human liver explants to historical data garnered from other preclinical models such as non-human primate and xenografted murine models. The comparison of AAV-LK03 and AAV8 might be the most relevant, given that clinical data for these vectors are publicly available in addition to data from multiple preclinical studies.^14, 15^ In the human explant under non-neutralizing conditions, AAV-LK03 surpassed AAV8 in both vector uptake and functional transduction (**Fig. 2d-e**). This aligns with cumulative clinical data for both capsids^14, 15^, however, it contradicts the hierarchical classification reported by Wang and colleagues in non-human primate studies^16^, where AAV8 outperformed AAV-LK03. Interestingly, Li and colleagues reported that rAAV3b, which only differs by 8 amino acids from AAV-LK03, substantially outperformed rAAV8 in transducing NHP livers.^17^ Our data are in line with those reported by Li *et al*. Specifically, when co-injected into a Cynomolgus monkey, both AAV-LK03 and the AAV3-like AAV-SEQ3 outperformed AAV8 at the level of vector uptake and transgene expression (**Fig. 4b**).

Interestingly, AAV-SYD12, which was the best performing variant at the level of transduction of the human liver explant in the absence of NAbs (**Fig. 2**), showed suboptimal performance in our NHP study (**Fig. 4b**), yet excelled targeting NHP primary hepatocytes within the FRG model (**Fig. 4c**). These findings suggest that AAV-SYD12’s *in vivo* efficacy in NHP may be affected by organ competition, which is absent in the murine non-liver organs. Alternatively, or additionally, the performance could be influenced by the blockage of transduction in primates due to existing neutralizing antibodies (1:30 NAb titer to AAV9). To draw concrete conclusions, further investigations are warranted. Lastly, it should be noted that the efficacy of AAV-SYD12 in the FRG model engrafted with NHP hepatocytes may be artificially overestimated, which is another potential explanation for the observed performance differences across the various models.

It is our opinion that the studies performed in the xenograft FRG mice model highlight the impact that the level of human hepatocyte engraftment can have on the outcome of experiments. Data obtained with AAV-LK03 provide for a good exemplar to illustrate this point. AAV-LK03’s performance varied depending on the degree of humanization, due to its high affinity for heparan sulfate proteoglycan (HSPG), a property contributing to a distinct periportal transduction profile.^18^ In low engrafted FRG mice, AAV-LK03 underperformed as fewer human hepatocyte clusters resided within the periportal zone. On the other hand, the relative performance of this capsid increased in highly engrafted mice where more human cells were randomly present within this zone. Using the human liver explant model, where all hepatocytes around the portal vein (AAVs entry route into the liver) are human, we found AAV-LK03’s performance closely matched that observed in highly engrafted mice. Therefore, when evaluating AAV vectors’ ability to transduce primary human hepatocytes, the use of highly engrafted animals could potentially ensure more accurate results.

It is important to state that the authors are not advocating replacement of the more tractable humanized FRG model with the whole human liver model. The FRG model has proven to be a robust preclinical tool, demonstrated by the successful development of vectors such as the AAV-SYDs and AAV-LK03 through directed evolution. However, we noted high discrepancies in transduction between the hFRG and human liver explant models for AAV-hu.Lvr06^19^ and AAV-LK03-REDH.^18^ Despite performing well in the humanized mouse model, these variants did not achieve the same level of performance in the liver explant model. This is specially intriguing for AAV-hu.Lvr06, a variant that was isolated directly from a human liver.^19^ It is possible that the low HSPG attachment of AAV-hu.Lvr06 and AAV-LK03-REDH could free up more vectors for human cell uptake in the vhFRG model, thereby artificially enhancing their bioavailability and overall relative performance. In the liver explant model, our data suggest these variants stay longer in the perfusate and performed better in the left graft of donor 1 where AAV was recirculated, providing more time for vector-cell interaction. In support of this, AAV-hu.Lvr06 outperformed all other variants in highly engrafted animals at two months post-injection, implying slower cell attachment/entry kinetics. Here it is important to note that our studies on vector performance in human livers explants were restricted to a roughly one-week timeframe. Given the observed dynamic nature of transgene expression observed in the FRG model, it is plausible that the relative functional transduction of the studied vectors, particularly at the RNA level, could exhibit variations with prolonged time frames. Therefore, while our data provides valuable insights into early vector performance and tissue tropism, future studies that examine these processes over more extended timeframes could further enrich our understanding of AAV vector behaviour and refine their use in gene therapy applications.

Finally, the *ex situ* human liver explant model introduces an exciting new avenue for the development of novel AAV vectors through directed evolution approaches. Its unique feature of maintaining native liver structure and human extracellular matrices makes it an ideal platform for refining strategies such as the HSPG de-targeting described previously in the humanized FRG model.^18^

All in all, we believe the human liver explant is among the most biologically and clinically predictive preclinical models available today, since it provides the valuable opportunity to perform vector development and evaluation in the context of the whole organ with intact organ architecture and zonation, in the presence of human blood. Further work needs to be conducted to understand convergences and divergences between the available preclinical models and how these relate to the accumulated clinical data. It is essential to highlight that this model is complementary to other *in vivo* models, such as the FRG or NHP, which continue to offer invaluable insights into vector biodistribution and AAV-mediated cellular toxicity.

In conclusion, we believe the whole human liver explant model offers the unprecedented opportunity to study AAV vector function in a model that closely recapitulates the human conditions. This novel model, with time, has the potential to lead to the development of the next-generation human liver targeted (and human liver de-targeted) AAV variants, starting a whole new era of potentially successful gene therapy liver directed clinical trials.

## Materials and Methods

### AAV transgene constructs

All the vectors used in the study contain AAV2 ITR sequences. The 44-mer long barcodes were cloned using standard molecular biological techniques into a self-complementary, CAG plasmid driving eGFP expression (Addgene #8327941). The sequences of such barcodes can be found in **Supplementary Table 2**. The barcodes were cloned between the fluorophore and the Simian virus 40 (SV40) polyadenylation as indicated in **Fig. 1b**.

### AAV vector packaging and viral production

AAV constructs were packaged into AAV capsids using human embryonic kidney (HEK) 293T cells and a helper-virus–free system as previously described. All AAV capsids used in this study were produced using polyethylenimine (PEI) transfection of the self-complementary CAG-eGFP-barcode transgene cassette, as well as adenovirus and Rep2-Cap helper plasmids using previously described methods.^20^ Briefly, each capsid serotype was co-transfected individually with the corresponding unique barcoded transgene. The resulting cell lysates were tittered individually and subsequently mixed at equimolar ratio. The vector mix was then purified using Cesium Chloride (CsCl) gradient ultracentrifugation following the previously published protocol.^21^

### AAV titration

AAV titration was performed via digital droplet PCR (ddPCR, Bio-Rad, Berkeley, CA, USA) using EvaGreen supermix (Bio-Rad, catalog no. 1864034) and following the manufacturer’s instructions. To detect AAV genomes on vectors, GFP primers were used (GFP-F: 5’-TCAAGATCCGCCACAACATC; GFP-R: 5’-TTCTCGTTGGGGTCTTTGCT).

### Barcode Amplification, NGS, and distribution analysis

To amplify the transgene region containing the 44-mer barcodes, BC-F (5’-GAGTTCGTGACCGCCG) and BC-R (‘5-ATTGCAGCTTATAATGGTTACAAATAAAGC) were used. NGS library preparations and sequencing using 2 × 150 paired-end configurations were performed by Azenta (Suzhou, China) using an Illumina HiSeq instrument. A workflow was written in Snakemake (5.6)42 to process reads and count barcodes. Paired reads were merged using BBMerge and then filtered for reads of the expected length in a second pass through BBDuk, both from BBTools 38.68 (https://sourceforge.net/projects/bbmap/). The merged, filtered fastq files were passed to a Python (3.7) script that identified barcodes corresponding to AAV variants. NGS reads from the DNA and cDNA populations were normalized to the reads from the pre-injection, vector mix.

### RNA stability test of the barcoded constructs

To minimize a possible bias in transgene RNA stability introduced by the longer barcodes, the 25 individual barcoded constructs were packaged in AAV2 and those constructs leading to either overexpression or under expression of the transgene when compared to the mean transduction values were discarded (**Supplementary Fig. 1**). Briefly, the 25 barcoded transgenes were mixed at 1:1 molar ratio and co-transfected into HEK293T as described below. After vector purification, HuH-7 cells were transduced as described previously with no modifications, with a multiplicity of infection of 5,000 vector genomes per cell. Three days after transduction, cells were harvested and the barcoded region was PCR-amplified from the vector preparation, DNA and RNA extracted from the cells, as described below.

### Cell Culture, Vector Transduction, and Heparin and Enoxaparin (Clexane) Competition Assay

HuH-7 cells were kindly provided by Dr Jerome Laurence (The University of Sydney). HEK293T cells were obtained from ATCC (Cat#CRL-3216). All cells were tested for mycoplasma and were mycoplasma-free. Cells were cultured as described previously (32490035) with no further modifications. For the heparin competition assay (**Supplementary Fig. 19**), cells were seeded at 10^5^ per well into 24-well plates at day 0 and transduced at the indicated vector genome/cell. When indicated, heparin sodium salt (Sigma, H3149-50KU, lot no. SLBW2119) or Clexane were supplemented at 50 μg/mL or at 200 μg/mL. After 72 h, the cells were harvested using TrypLE express and analyzed for GFP using BD LSRFortessa cell analyzer. The data were analyzed using FlowJo 7.6.1.

### Enzyme-linked Immunosorbent Assay (ELISA) measurement of anti-AAV IgG specific antibody titer in human serum

Human sera were assayed for reactivity to all the fourteen capsids by ELISA, following a recently described method.^8^ 96-well polystyrene ELISA plates (Nunc #442404) were coated overnight at 4°C with 50 μL per well of the AAV vector mix (2.5×10^10^ vg/mL) diluted in coating buffer (carbonate-bicarbonate buffer, SigmaAldrich). Plates were washed 3 times with wash buffer (PBS) + 0.05% Tween-20 (Sigma Aldrich) and then received 100 μL per well of blocking buffer (PBS + 5% skim milk + 0.05% Tween-20). Plates were then washed three times after incubation at room temperature for 2 hours in wash buffer and received 50 μL per well of sera (diluted in blocking buffer at 1:50 and at 1:200 with duplicate wells for each dilution). Plates were incubated for 2 hours at room temperature and washed 3 times with wash buffer before receiving 50 μL per well of horse radish peroxidase (HRP)-conjugated anti-human Fc specific IgG (Chemicon AP309P, diluted 1:10,000 in blocking buffer). Plates were incubated for 1 hour at room temperature and washed 4 times using wash buffer before receiving 75 μL per well of 3,3′,5,5′-Tetramethylbenzidine (TMB, Sigma-Aldrich). Plates were incubated in the dark for 30 minutes at room temperature and the reactions were then stopped using 75 μL per well of 1M sulfuric acid. The absorbance of each well was measured at 450 nm wavelength using a VersaMax microplate reader (Molecular Devices, LLC). Duplicate wells containing no AAV served as background controls. The mean value for each sample dilution was calculated for wells with (foreground) and without coated vector (background) and the sample was considered reactive if this ratio was >2.0. Reactive and non-reactive human sera to the whole mix were subsequently assayed for reactivity to all the fourteen capsids individually following the same protocol. The chosen sera were diluted in blocking buffer at 1:25, 1:50, and at 1:100 with duplicate wells for each dilution.

### Liver origin and procurement

The two livers described in this manuscript were obtained from DonateLife, the centralized donation organization in Australia. These livers were unsuitable for transplantation but consented for research use. The first liver was medically unsuitable for transplantation due to biliary sepsis and cholecystostomy, whereas the latter was a donation after circulatory death (DCDD) that did not meet the acceptance criteria for transplantation of age <50. Livers were procured in our standard fashion with aortic flushing using cold Soltran (Baxter Healthcare, Illinois, USA) and University of Wisconsin preservation solution (Belzer UW, Bridge to Life, Columbia, USA). In addition, DCDD livers received an intra-aortic injection of tissue plasminogen activator (Alteplase 20mg) delivered as a bolus at the time of cold perfusion. All livers were placed in static cold storage for transfer to our center.

### Liver viability test

At the indicated time points, we assessed both human livers for viability, following the criteria proposed by the VITTAL clinical trial.^3^ These are a concentration of lactate ≤2.5mmol/L, and two or more of: bile production, pH≥7.30, glucose metabolism, hepatic arterial flow ≥150ml/minute and portal vein flow ≥500ml/minute, or homogeneous perfusion.

### Liver assist modifications

We utilised the recently described^5^ modified commercial liver perfusion system (Liver assist, Organ assist, Gronigen, Netherlands), as we have previously described which uses an open venous reservoir. Briefly, we added a flow adjustable dialysis membrane (Prismaflex or Polyflux, Baxter Healthcare, Illinois, USA) for filtration of water soluble toxins and control of perfusate volume, long term oxygenators (Quadrox-iD Pediatric, Macquet, Getinge Group, Rastatt, Germany) for extended perfusion, and a gas blender (Device Technologies, Sydney, Australia) with a paediatric flow regulator for fine control of ventilation. The perfusate contained 4 units of human packed red cells, 2 units of fresh frozen plasma, 200mL of 20% albumin and 1L of normal saline. Anticoagulation was maintained using enoxaparin (100mg twice daily) and nutritional support provided using infusions of amino acids (Synthamin 17, Baxter Healthcare, Illinois, USA), lipids (Clinoleic 20%, Baxter Healthcare, Illinois, USA), taurocholic acid (7.7 mg/h), methylprednisolone (21mg/h), insulin (Actrapid 2 IU/ml) and glucagon (20 ug/mL).

Prior to connecting the liver to the perfusion machine, we circulated the perfusate and activated the parallel dialysis circuit until the potassium, calcium and acidosis abnormalities in the stored blood were corrected. After flushing with saline, we connected the liver to the system by portal vein and hepatic artery cannulas and we rewarmed the perfusate at 1 degree/h from 32°C to 36°C to minimise haemolysis.

### Liver splitting

The *ex situ* split was performed the day after commencement of whole liver perfusion, which corresponded to a whole liver perfusion time of 12-16 hours. Splitting into a left lateral sector graft (LLSG, segments 2 and 3) and an extended right graft (ERG, segments 1 and 4-8) was performed without interrupting arterial or portovenous perfusion and required 2 surgeons and a scrubbed assistant. The procedure itself was divided into 3 phases which have been described in detail recently.^22^

### Measurements of liver function

Liver synthetic function was assessed by measuring lactate clearance, bile production and perfusate biochemistry. We measured lactate concentration, pH, glucose concentration, pO_2_ and pCO_2_ with a blood gas analyser (RAPIDPoint 500, Siemens Healthengineers, Norwood, Massachusetts, USA). The liver function tests, coagulation studies, and Factor V were measured by the clinical laboratory of the Royal Prince Alfred Hospital, Sydney, Australia. We assessed the architectural integrity of the liver by staining formalin-fixed core biopsies with haematoxylin and eosin. Biopsies were also scored by a specialist pathologist for percentage of coagulative necrosis, hepatocyte detachment, bile duct injury (**Supplementary Figs 20, 21, and 22**).

### DNA and RNA isolation and cDNA synthesis

Isolation of DNA, RNA, and cDNA synthesis was performed as described in detail before^19^ with no further modifications. Briefly, DNA was extracted using a standard phenol: chloroform protocol and RNA with the Direct-Zol kit (Zymogen Cat #R2062). cDNA synthesis was carried out with SuperScript IV first-strand synthesis (Invitrogen), following manufacturer’s instructions.

### Immunofluorescence analyses

Immunofluorescence was performed, as described recently in detail recently without modifications.^6^ Briefly, liver tissue was fixed with paraformaldehyde, cryo-protected in sucrose, and frozen in O.C.T. (Tissue-Tek). Frozen liver sections (5 μm) were permeabilized in ice-cold methanol, then room temperature 0.1% Triton X-100, and then reacted with DAPI at 0.08 ng/mL (Invitrogen, D1306) and the antibodies indicated below. Anti-eGFP (Invitrogen, #A-11122), Anti-HNF4α (Abcam, #ab41898), Anti-HAL (Sigma Prestige Antibodies, #HPA038547), anti-Cytokeratin 7 (Abcam, #ab68459), anti-albumin (Bethyl, #A80229A), anti-KIAA0319L (AAVR, Abcam, #ab105385), anti-CD68 (Invitrogen, #14068882) antibodies were used as indicated. For **Fig. 2-k**, and **3h-k** antigen retrieval was required. This was achieved by heating the slides for ten minutes in antigen retrieval buffer (Tris-EDTA, pH 9). After immunolabelling, the images were captured and analyzed on a LSM800-Airyscan microscope using ZEN Black software.

### Mouse studies and isolation of human hepatocytes by collagenase perfusion

All animal experimental procedures and care were approved by the joint Children’s Medical Research Institute (CMRI) and The Children’s Hospital at Westmead Animal Care and Ethics Committee. Fah^−/-^Rag2^−/−^Il2rg^−/-^ (FRG) mice were bred, housed, engrafted, and monitored as recently described.^6^ Levels of human and non-human primate cell engraftment were estimated by measuring the presence of human albumin in peripheral blood, using the human albumin ELISA quantitation kit (Bethyl Laboratories, #E80-129). To evaluate the AAV transduction potential, mice were placed on 10% NTBC and were maintained in this condition until harvest. All hFRG mice reported in this study were engrafted with human and non-human primate hepatocytes from the same donors (Caucasian, 15-month-old donor, Lonza, #HUM181791; ThermoFisher, #CY409, respectively).

Mice were randomly assigned to experiments and transduced via intravenous injection (lateral tail vein) with the indicated vector doses. Mice were euthanized by CO_2_ inhalation either 1 week after transduction for barcoded NGS analyses, unless indicated otherwise. To obtain murine and human/non-human primate single-cell suspensions from xenografted murine livers, we followed the same collagenase perfusion procedure as recently described.^6^ Briefly, cells were labelled with phycoerythrin (PE)-conjugated anti-human-HLA-ABC (clone W6/32, Invitrogen 12-9983-42; 1:20), biotin-conjugated anti-mouse-H-2Kb (clone AF6-88.5, BD Pharmigen 553,568; 1:100), and allophycocyanin (APC)-conjugated streptavidin (eBioscience 17-4317-82; 1:500). Flow cytometry was performed in the Flow Cytometry Facility, Westmead Institute for Medical Research (WMIR), Westmead, NSW, Australia. The data were analyzed using FlowJo 7.6.1 (FlowJo LLC).

### Non-human primate work

Animal procedures were approved by the ethical committee for animal testing of the University of Navarra and by the Department of Health of the government of Navarra (Comité de Etica para la Experimentación Animal code: 038/15) and performed according to the guidelines from the institutional ethics commission. Animal welfare checks were performed by animal care staff twice daily. A young adult male Macaca fascicularis NHP animal was subjected to the immunoadsorption process (described in detail in ^23^). Within the following 30 min after immunoadsorption, the vector was infused via the saphenous vein over ten minutes. At day 30, the animal was euthanized, and the four liver lobes were collected for further analysis.

## Acknowledgements

We thank the Cytometry Facility of the Westmead Institute for Medical Research for help with sorting of human/murine hepatocytes. We also would like to thank all the members of the Children’s Medical Research Institute Bioresources facility, with special thanks to S. Dimech. We would also like to thank Dr Razvan Albu for help generating the barcoded transgenes, Dr Anais Karime Amaya for initial assessment of liver explant viability, and Dr Grant Logan for his guidance on the ELISA procedure described herein.

This work was supported by project grants from the Australian National Health and Medical Research Council (NHMRC) to L.L. (APP2021305 and APP1161583). The work of L.L. was also supported by a research grant from the National Science Centre, Republic of Poland (OPUS-21) (2021/41/B/NZ5/01671).

M.C-C. was also supported by a 2021 New South Wales (NSW) Ministry of Health, Office of Health and Medical Research (OHMR) Early-Mid Career Research Grant - Gene and Cell Therapy. Financial support was also provided by the Royal Prince Alfred Hospital Transplant Institute. N-G.L is supported by the Australian Government Research Training Program Stipend Scholarship.

## Author contributions

Conceptualization, M.C-C., I.E.A., and L.L.; methodology, M.C.-C., N-S. L., M.L., C.U., G.G-A, and C.P.; investigation: M.C.-C., S.H.Y.L., R.G.N., M.K., D.N., N-S.L., M.L., E.Z., R.B.D., A.W., and G.B.; writing – original draft, M.C.-C.; writing – review and editing, M.C.-C., I.E.A., and L.L.; funding acquisition, M.C-C., G.M., C.U., G.G-A, I.E.A., C.P., and L.L.; visualization, M.C.-C., J.M., R.R.-P.; supervision, M.C.-C., G.M., C.U., G.G-A, I.E.A., C.P., and L.L.

