## Supplemental Figures for "Harnessing the power of whole human liver ex situ normothermic perfusion for preclinical AAV vector evaluation"

### Supplementary Table 1. AAV vectors used in the study.

| AAV variants | Origin | Method | Composition | Reference |
|---|---|---|---|---|
| AAV5 | Nature - Human | Isolation | Wild-type | 6324476 |
| AAV8 | Nature - NHP | Isolation | Wild-type | PMC129358 |
| AAV9 | Nature - Human | Isolation | Wild-type | PMC416542 |
| AAV-hu.Lvr06 | Nature - Human | Isolation | Wild-type – AAV2-like | 32908003 |
| AAV-LK03 | Bioengineered | Directed Evolution - hFRG | Shuffled – AAV3b-like | 24390344 |
| AAV-SEQ3 | Bioengineered | Empirical domain swap | Domain Swap – AAV3b-like | WO/2016/181123 |
| AAV-FT01 | Bioengineered | Directed Evolution - hFRG | Peptide Displayed on AAV2 | 35297686 |
| AAV-FT11 | Bioengineered | Directed Evolution - hFRG | Peptide Displayed on AAV2 | 35297686 |
| AAV-SYD11 | Bioengineered | Directed Evolution - hFRG | Shuffled – AAV7-like | 34977275 |
| AAV-SYD12 | Bioengineered | Directed Evolution - hFRG | Shuffled – AAV7-like | 34977275 |
| LK03-REDH | Bioengineered | Rational Design | R594E+D598H | 36700121 |
| AAV8-Swap03 | Bioengineered | Domain Swap - hFRG | AAV8 with VR-VI to VIII from AAV7 | 34977275 |
| AAV8-Swap05 | Bioengineered | Domain Swap - hFRG | AAV8 with VRI from AAV2 + VR-VI to VIII from AAV7 | 34977275 |
| AAV8-Swap06 | Bioengineered | Domain Swap - hFRG | AAV8 with VR-IV and V from AAV10 + VRVI-VIII AAV7 | 34977275 |

### Supplementary Table 2. Barcode Sequences.

| Barcode | Capsid | Sequence (5' to 3') |
|---|---|---|
| BC2 | AAV-hu.Lvr06 | GATAGGGAAGCAAAGCAAACTAAACTGAACGGAACGAAGGGGAA |
| BC5 | AAV-SEQ3 | GGTCGCTTGGTGTAGGTACGGTCACGACAGTGAACCTCGGCATC |
| BC9 | AAV5 | TGTAGGGAGGTGTACTGTCCCGAACCGTAGGGAAGTGCGGGAAC |
| BC10 | AAV-SYD11 | TTCGCTTAGCTATGCCTCAGTTAACGACCCTGCACTTCAGCACC |
| BC11 | AAV-SYD12 | TGAACCTTAGCGAGCTACCGTATGGCATGGGGAACCATCCGACC |
| BC12 | AAV8-Swap03 | TTAACTTAAGCGCACCACAGGATAGCAAAGTTCACCAAACTTAA |
| BC13 | AAV8-Swap05 | TACACGGTCCTTTGGGACACTGCGGTACAGCAAAGCGTCCTACC |
| BC14 | AAV8-Swap06 | TTTGGTGTGCGTCCCCAAACCAAACTAACCGGCGCCTTACGACA |
| BC15 | AAV-FT01 | CTAACTGAAGTACCCGACGCCGCCGGGAGCTTTACTGAACCTAA |
| BC18 | AAV-FT11 | CGAAGCTAGGTGCACCAAACCTTGCGTAACCACGCTGTGGCAAA |
| BC20 | AAV-LK03 | TACACCTAAGTGACCTTTCCCTAAGGTCAGCGACGTTTGGTTAA |
| BC22 | LK03-REDH | CTTCGTTAAGCTCCGCGACCCTAACTGCACGGAACGATAGGAAA |
| BC23 | AAV8 | GTAGGGAAGCGATGCCTACCGGAGGGAAGGCGCACCAACGTTCA |
| BC24 | AAV9 | TGTACCAAACGGAAGGGAATCTTGCCACACTACGCCAACCGGTG |

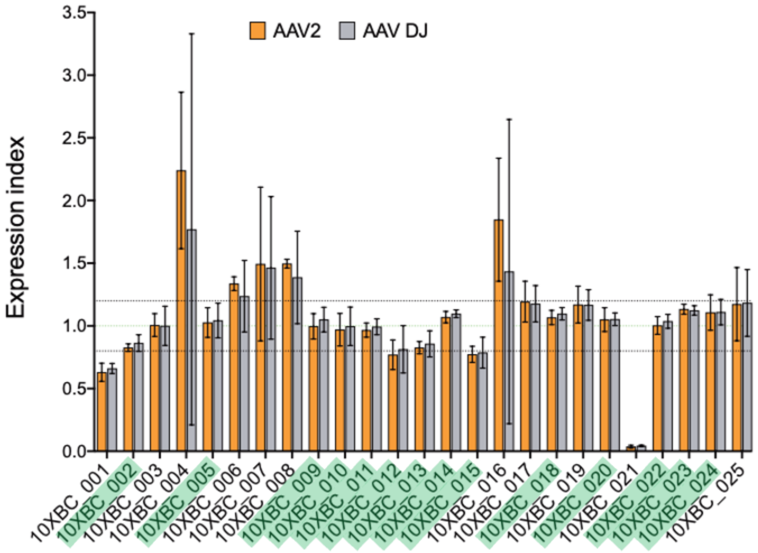


**Supplementary Figure 1. RNA Stability test of the barcoded transgenes.** To minimize a possible bias in transgene RNA stability introduced by the longer barcodes, the 25 individual barcoded constructs were packaged in AAV2 and AAV-DJ, and those constructs leading to either overexpression or under expression of the transgene when compared to the mean transduction values were discarded. Briefly, the 25 barcoded transgenes we were mixed at 1:1 molar ratio and co-transfected into HEK293T as described in the **Material and Methods** section for vector production. After AAV2 and AAV-DJ vector purification, 293T cells were transduced at three multiplicities of transduction (50, 500, and 5000 vector genomes per cell). Three days after transduction, cells were harvested and the barcoded region was PCR-amplified from the vector preparation, DNA and RNA extracted from the cells. The results presented herein show the quotient of NGS reads from mRNA and gDNA. N=3 multiplicities of transduction. The green highlighted barcodes were subsequently packaged in each capsid described in **Supplementary Table 2**.

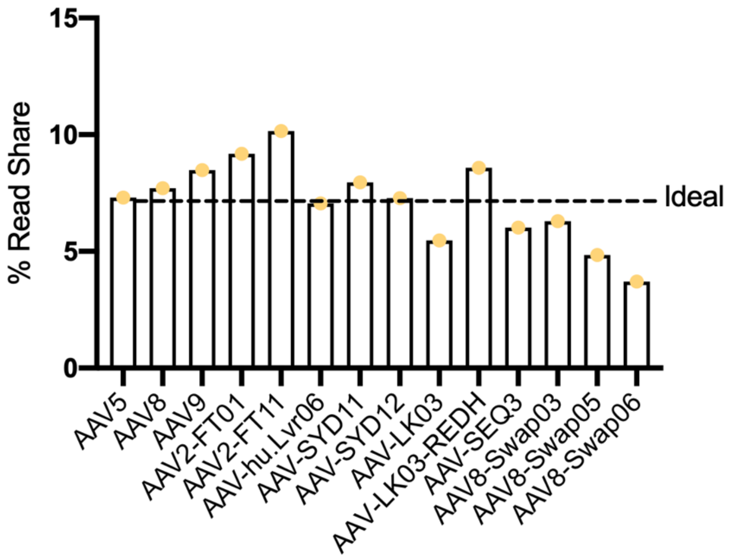


**Supplementary Figure 2. Barcoded transgene pre-mix validation.** Results show the percentage of NGS-reads mapped to each barcoded transgene packaged in the displayed AAV variants. The transgene was amplified with PCR from the vector preparation. Given that fourteen capsids were present in the mix, an ideal distribution would contain 7.14% of each variant, showed with a dashed line. All the results obtained throughout the manuscript, unless indicated otherwise, were normalized to this vector pre-mix dataset.

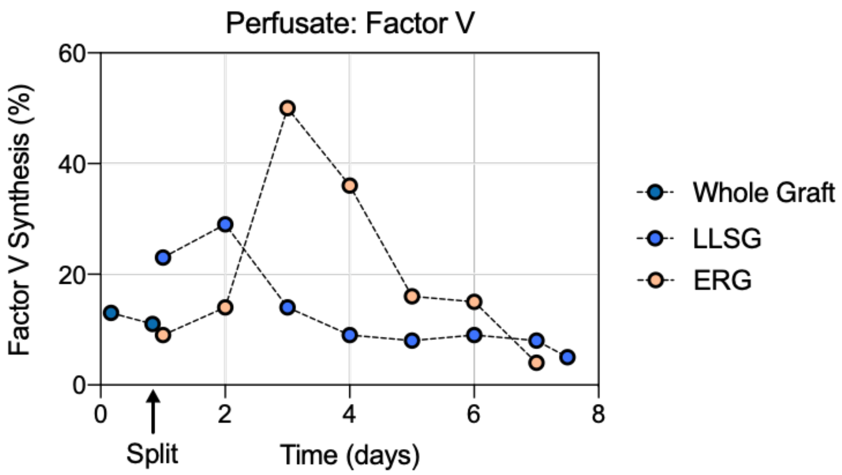


**Supplementary Figure 3. Factor V synthesis.** Course of Factor V synthesis for Donor 1, expressed as a percentage of the normal value, as studied *a posteriori* in samples taken from the perfusate throughout the perfusion. LLSG – Left Lateral Sector Graft. EG – Extended Right Graft.

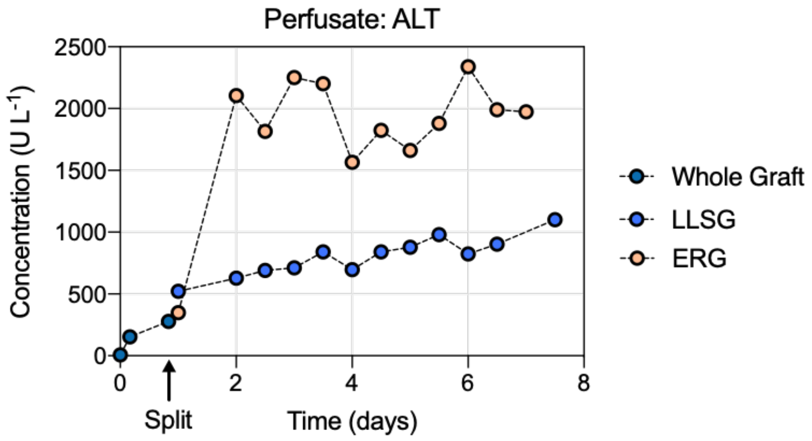


**Supplementary Figure 4. Alanine aminotransferase (ALT) levels.** Course of Alanine aminotransferase concentration in the perfusate for Donor 1, as studied *a posteriori* in samples taken from the perfusate throughout the perfusion. LLSG – Left Lateral Sector Graft. EG – Extended Right Graft.

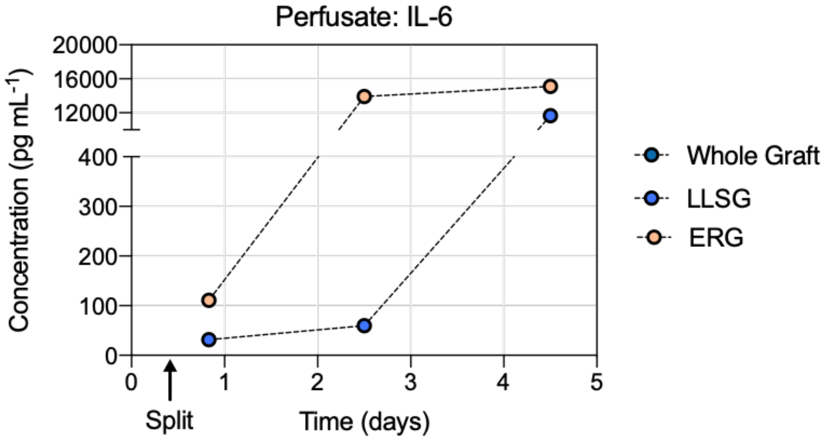


**Supplementary Figure 5. Interleukin 6 (IL-6) levels.** Course of IL-6 concentration in the perfusate for Donor 1, as studied *a posteriori* in samples taken from the perfusate throughout the perfusion. LLSG – Left Lateral Sector Graft. EG – Extended Right Graft.

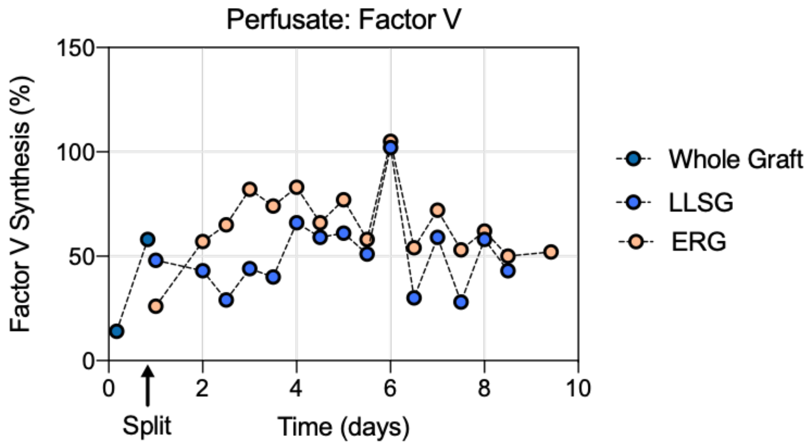


**Supplementary Figure 6. Factor V synthesis.** Course of Factor V synthesis for Donor 2, expressed as a percentage of the normal value, as studied *a posteriori* in samples taken from the perfusate throughout the perfusion. LLSG – Left Lateral Sector Graft. EG – Extended Right Graft.

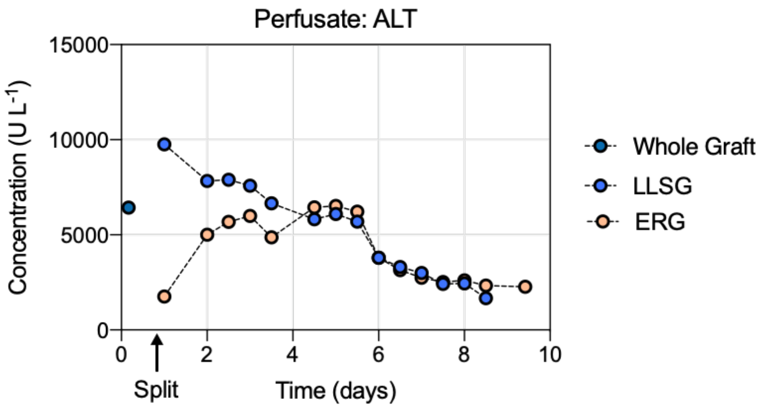


**Supplementary Figure 7. Alanine aminotransferase (ALT) levels.** Course of Alanine aminotransferase concentration in the perfusate for Donor 2, as studied *a posteriori* in samples taken from the perfusate throughout the perfusion. LLSG – Left Lateral Sector Graft. EG – Extended Right Graft.

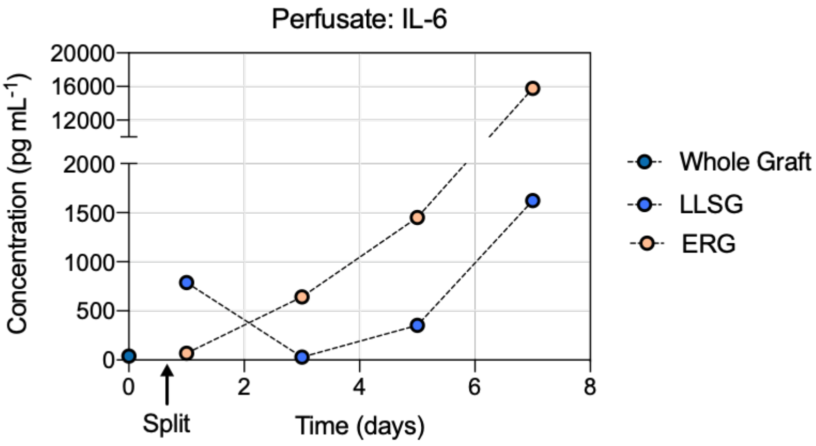


**Supplementary Figure 8. Interleukin 6 (IL-6) levels.** Course of IL-6 concentration in the perfusate for Donor 2, as studied *a posteriori* in samples taken from the perfusate throughout the perfusion. LLSG – Left Lateral Sector Graft. EG – Extended Right Graft.

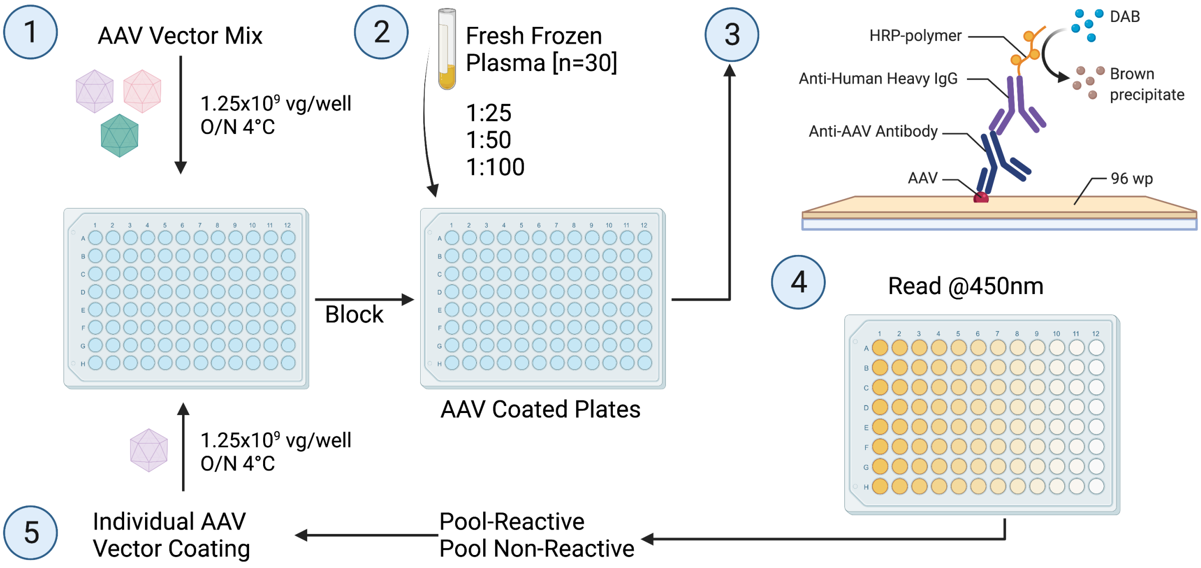


**Supplementary Figure 9. ELISA method outline.** Human sera were assayed for reactivity to all the fourteen capsids by ELISA, following a recently described method as outlined in the Materials and Methods section.

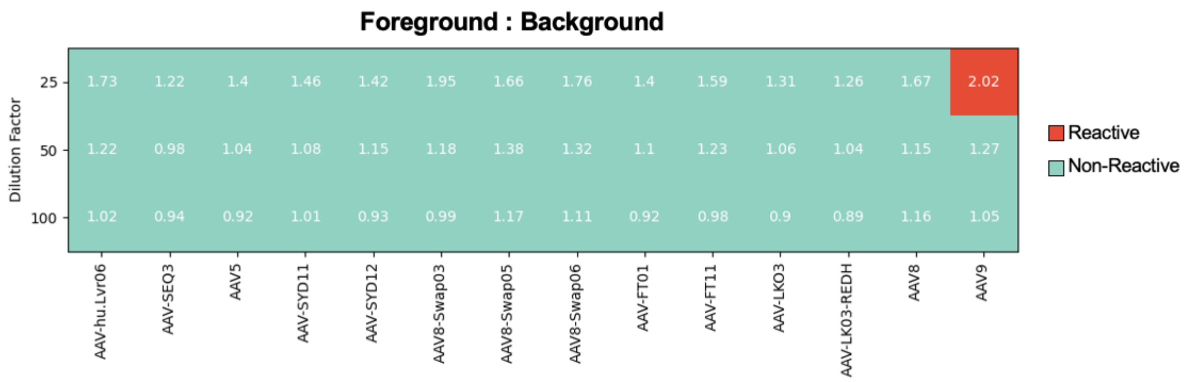


**Supplementary Figure 10. Reactivity of the 'non-reactive' human plasma used in the perfusion of Donor 1.** The mean value for each sample dilution was calculated for wells with (foreground) and without coated vector (background) and the sample was considered reactive if this ratio was >2.0. Only AAV9 was found to be slightly reactive at 1:25.

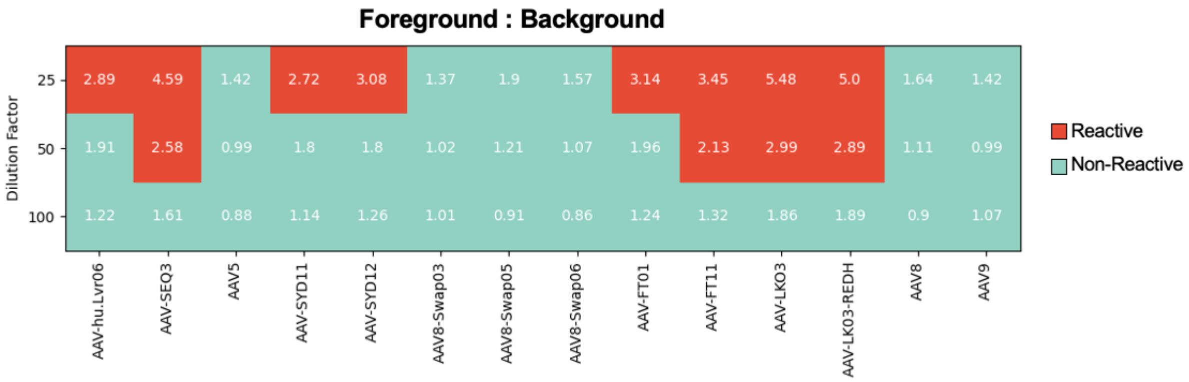


**Supplementary Figure 11. Reactivity of the 'reactive' human plasma used in the perfusion of Donor 2.** The mean value for each sample dilution was calculated for wells with (foreground) and without coated vector (background) and the sample was considered reactive if this ratio was >2.0.

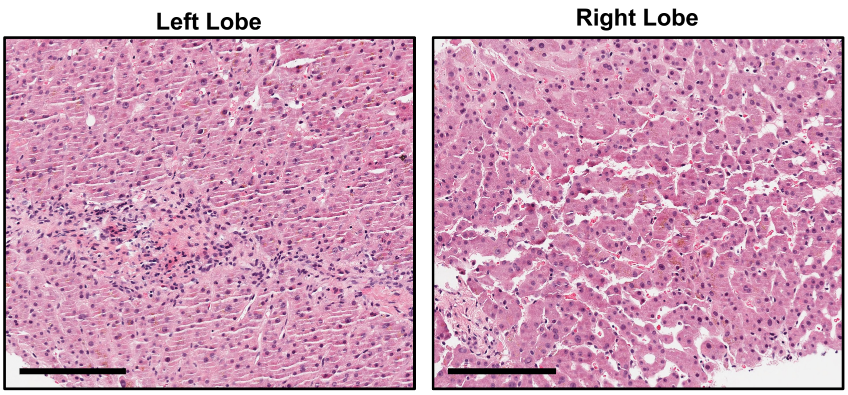


**Supplementary Figure 12. Liver histology of representative core biopsies from Donor 1.** Hematoxylin and eosin stain. Scale: 200 µm.

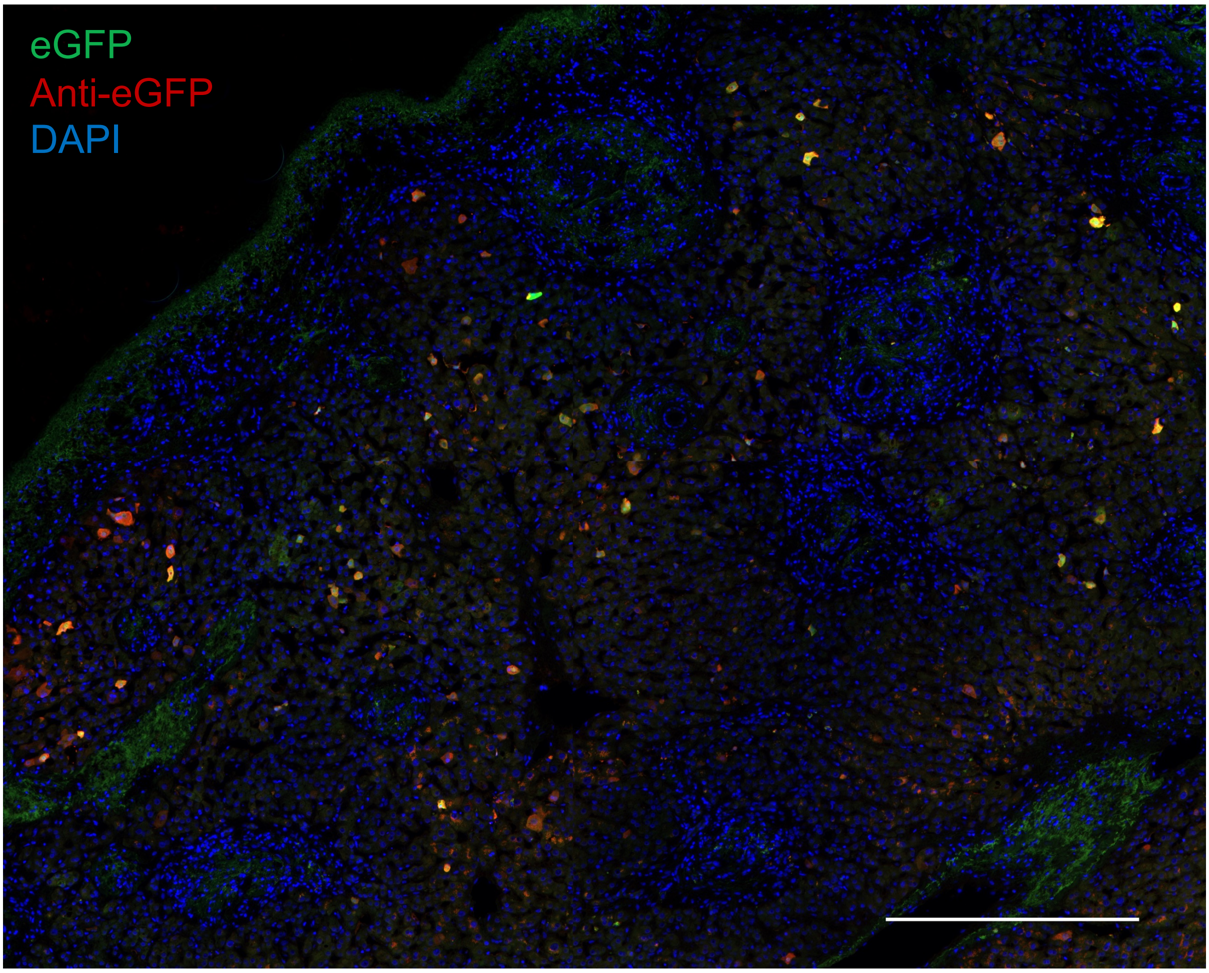


**Supplementary Figure 13. Net eGFP signal.** Representative immunofluorescence analysis of the net eGFP signal from collective AAV transduction in the left lobe of Donor 1 at day 4 post-transduction. The vector-encoded eGFP was also counterstained with an anti-eGFP antibody (red). Blue: DAPI (nuclei). Scale: 500 µm.

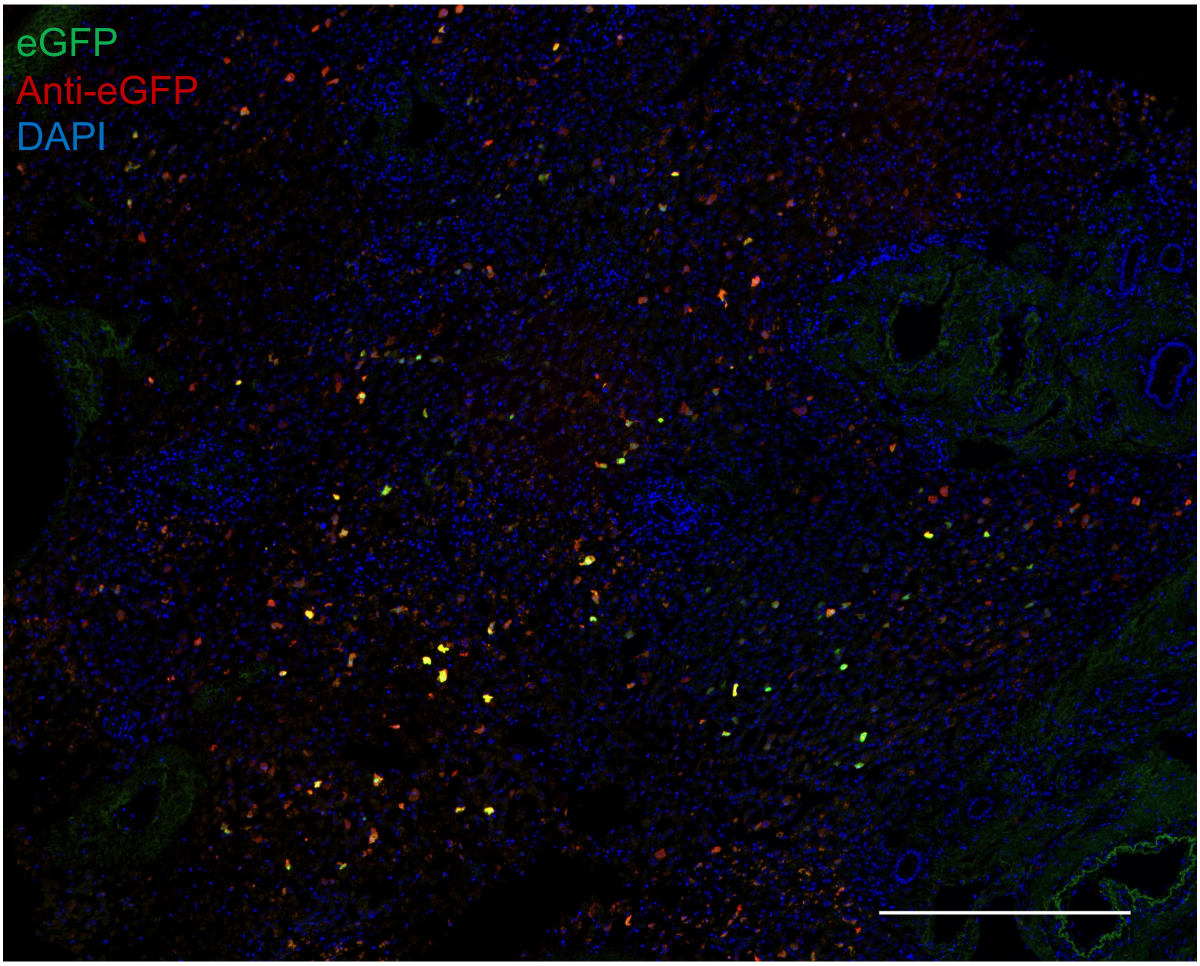


**Supplementary Figure 14. Net eGFP signal.** Representative immunofluorescence analysis of the net eGFP signal from collective AAV transduction in the right lobe of Donor 1 at day 4 post-transduction. The vector-encoded eGFP was also counterstained with an anti-eGFP antibody (red). Blue: DAPI (nuclei). Scale: 500 µm.

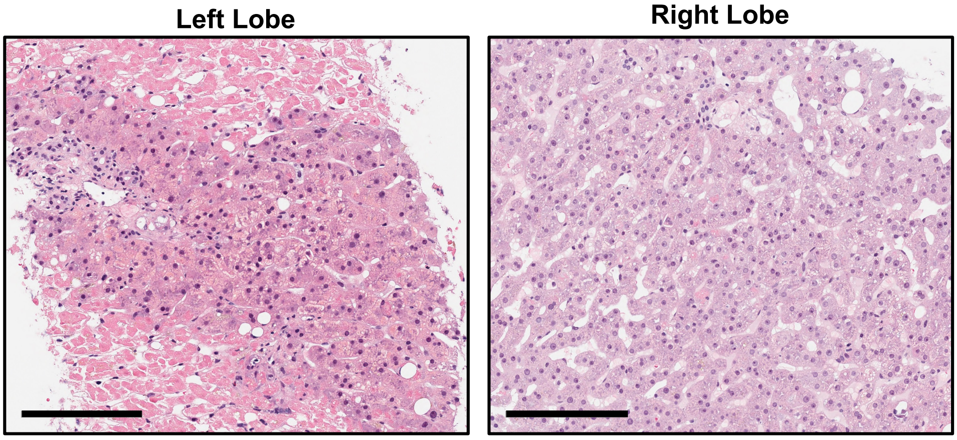


**Supplementary Figure 15. Liver histology of representative core biopsies from Donor 1.** Hematoxylin and eosin stain. Scale: 200 µm.

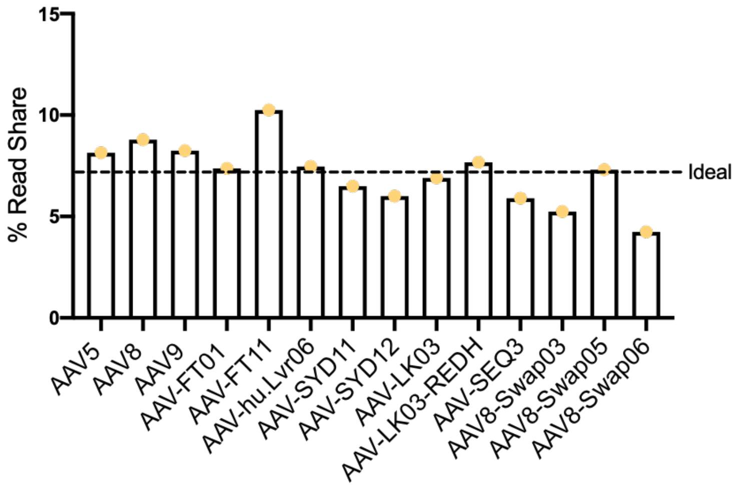


**Supplementary Figure 16.** Results show the percentage of NGS-reads mapped to each barcoded transgene packaged in the displayed AAV variants injected in the non-human primate described in **Fig. 4b**. The transgene was amplified with PCR from the vector preparation. Given that fourteen capsids were present in the mix, an ideal distribution would contain 7.14% of each variant, showed with a dashed line.

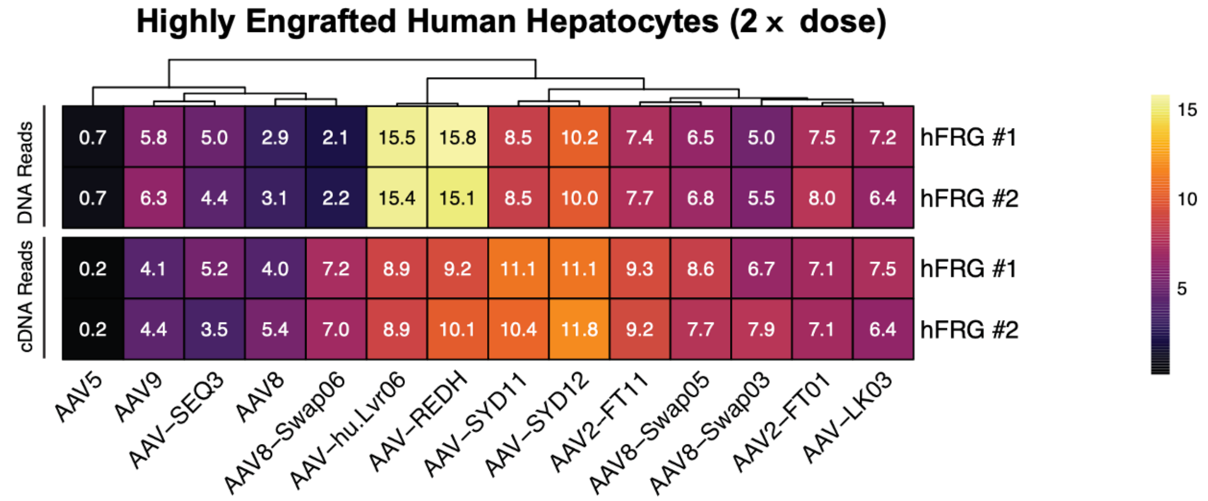


**Supplementary Figure 17. Transduction of highly humanized FRG mice with a total dose of 2.1×10$^{11}$ vector genomes.** Percentage of NGS reads mapped to each barcoded AAV capsid variant. On the top panel, the transgene DNA, indicating vector uptake, was extracted from sorted human hepatocytes, one-week post-injection. The bottom panel shows a similar analysis performed on transgenes recovered from RNA, which indicate functional transduction. Percentages are normalized to the pre-injection mix.

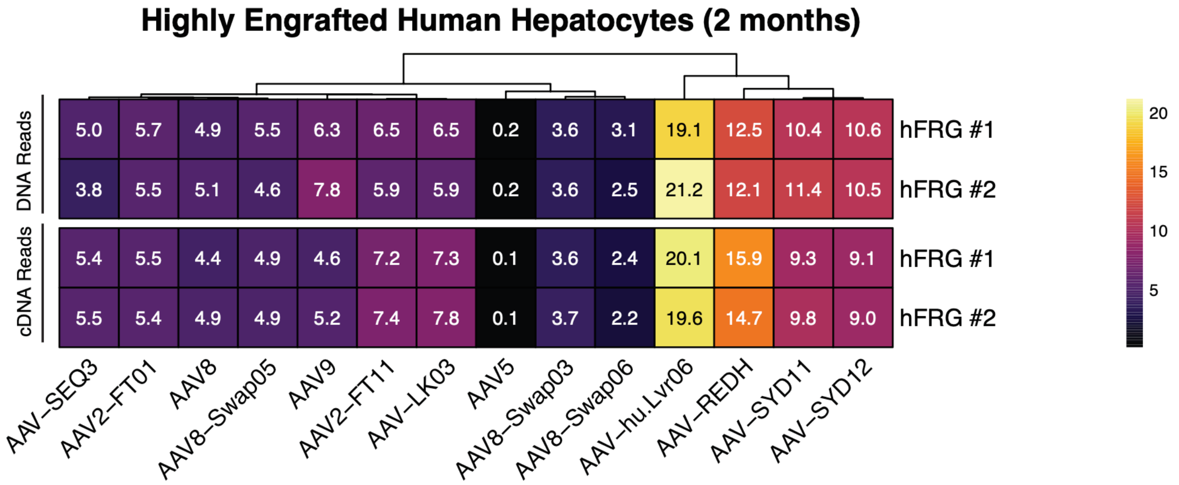


**Supplementary Figure 18. Transduction of highly humanized FRG mice as analyzed two months after injection.** Percentage of NGS reads mapped to each barcoded AAV capsid variant. On the top panel, the transgene DNA, indicating vector uptake, was extracted from sorted human hepatocytes, one-week post-injection. The bottom panel shows a similar analysis performed on transgenes recovered from RNA, which indicate functional transduction. Percentages are normalized to the pre-injection mix.

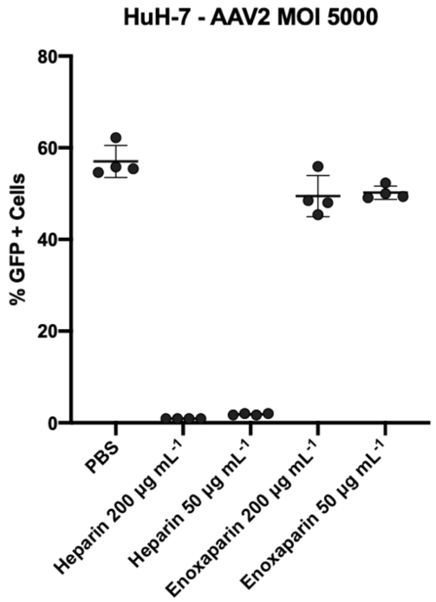


**Supplementary Figure 19. Heparin and enoxaparin competition assay.** Cells were transduced at 5000 vector genomes/cell. When indicated, heparin sodium salt or enoxaparin were supplemented at 50 or at 200 µg $mL^{-1}$. After 72 h, the cells were harvested and analyzed for GFP expression, as indicated on the y-axis

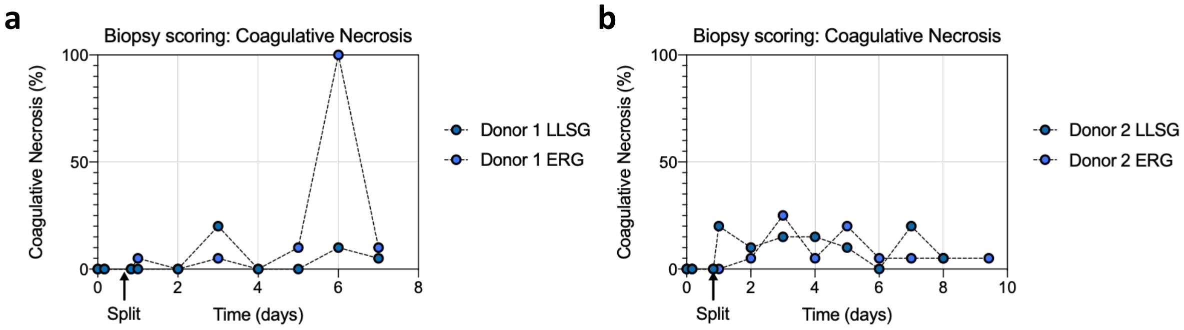


**Supplementary Figure 20. Coagulative necrosis of livers (%).** Core biopsies taken at the indicated time points were assessed by a specialist pathologist. **a,** Donor 1; **b,** Donor 2.

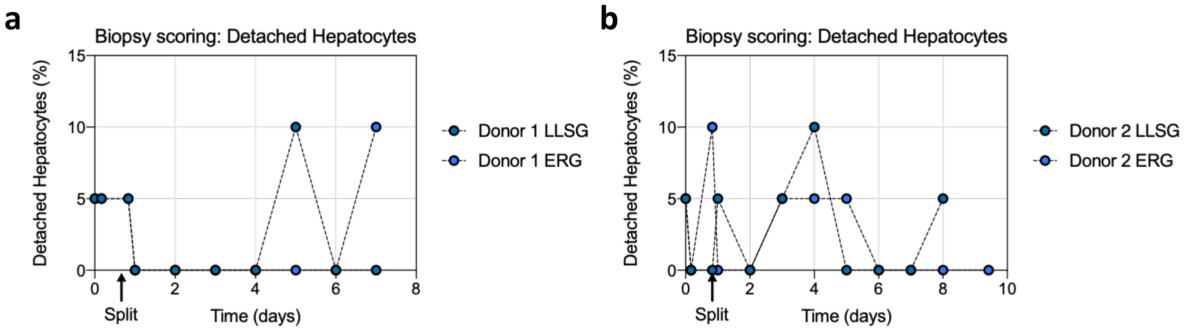


**Supplementary Figure 21. Detached hepatocytes (%).** Core biopsies taken at the indicated time points were assessed by a specialist pathologist. **a,** Donor 1; **b,** Donor 2.

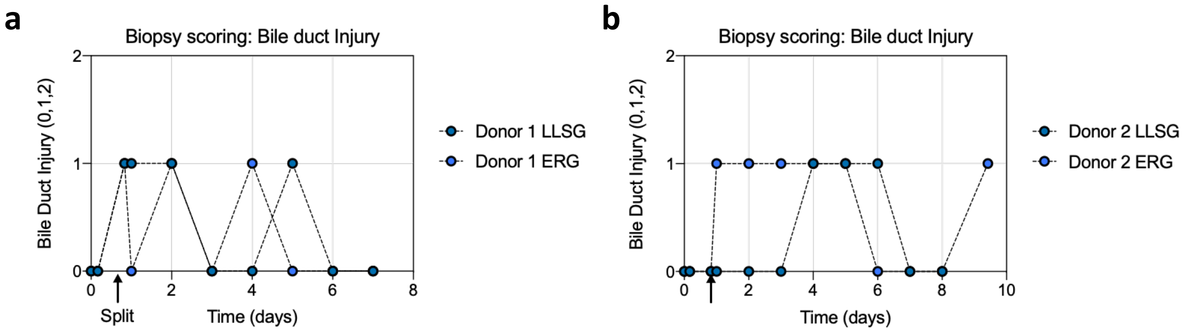


**Supplementary Figure 22. Bile duct Injury.** Core biopsies taken at the indicated time points were assessed by a specialist pathologist. **a,** Donor 1; **b,** Donor 2.
